# Cultured endothelial cells present organ-specific RTK distributions: advancing receptor measurement and data standardization via quantitative flow cytometry

**DOI:** 10.1101/2025.02.27.640662

**Authors:** Xinming Liu, Yingye Fang, Princess I. Imoukhuede

## Abstract

**Purpose:** Receptor tyrosine kinase (RTK) concentrations on the plasma membrane correlate with angiogenic functions in vitro and in rodent models. The intracellular RTK pool also regulates plasma membrane receptor availability and signaling pathways. Organs have specialized angiogenic functions essential to their distinct roles, supporting the hypothesis that plasma membrane and intracellular RTK concentrations vary across endothelial cells (ECs) from different organs.

**Methods:** Using quantitative flow cytometry on human ECs derived from dermis, umbilical vein, kidney, liver, and brain, we measured and statistically analyzed the concentrations of selected RTKs within ECs and on their plasma membranes.

**Results:** VEGFR1 exhibited the lowest concentrations on the plasma membrane (300–900 VEGFR1/cell) among VEGFRs. HDMECs (dermis) showed the lowest VEGFR1 level among the examined EC types. Whole-cell VEGFR1 concentrations were 2500–7500 VEGFR1/cell, with 12–26% located on the plasma membrane. The proportion of VEGFR2 located on the plasma membrane was higher at > 30%, except in HGMECs (kidney) where it was 24%. Plasma membrane VEGFR2 was significantly lower in HDMECs and HGMECs compared with HBMECs (brain), whereas whole-cell VEGFR2 levels were consistently in the range of 14,100–22,500 molecules/cell. VEGFR3 was the least localized to the plasma membrane, from 2% in HGMECs to 14% in HDMECs at the highest level of 4400 VEGFR3/cell. Whole-cell VEGFR3 concentrations ranged from 32,400 in HDMECs to 62,000 VEGFR3/cell in HLiSMECs (liver), with no significant differences among EC types. NRP1 was most abundant on the plasma membrane of HUVECs (umbilical vein) at 39,700 NRP1/cell; other ECs displayed 26,000–29,900 NRP1/cell, approximately 5-fold higher than the numbers of VEGFRs. Across EC types, Axl was present on the plasma membrane at levels (6900–12,200 Axl/cell) similar to those of VEGFR2.

**Conclusions:** We quantified and statistically analyzed plasma membrane and whole-cell expression of angiogenic RTKs across cultured human ECs from five different organs. Our findings suggest that RTK protein distribution might not fully reflect the differential angiogenic capacities in cultured ECs. In vitro monoculture conditions might reduce EC organ-specific features essential for refining vascular models.

## Introduction

Endothelial cells (ECs) lining blood vessels are not uniform across all tissues; they exhibit a range of differences, known as phenotypic heterogeneity. This diversity can be traced back to the earliest stages of vascular development, which begins with the differentiation of endothelial cells (ECs) and the formation of a primitive vascular network [1]. As development progresses, the network remodels and acquires organ-specific functions in response to alterations in hemodynamics and the surrounding microenvironment [1]. Initially homogeneous ECs differentiate into functionally distinct cells with features tailored specifically for the endothelia of various organs [2]. Investigating this phenotypic heterogeneity can uncover how specific EC types uniquely support the health and function of each organ.

Angiogenesis is regulated mostly via transmembrane receptor tyrosine kinases (RTKs), including vascular endothelial growth factor receptors (VEGFRs) and their coreceptor neuropilins (NRPs), Axl, and platelet-derived growth factor receptors (PDGFRs) [3]. These receptors on the plasma membrane bind growth factor ligands and transduce pro- or anti-angiogenic signals into intracellular compartments [3]. Because vascular diseases are usually associated with dysregulated RTK expressions, it is essential to characterize RTK densities on ECs and explore their association with function and their possible utility as biomarkers of vascular diseases [4].

Studies have measured the abundance of plasma membrane RTKs in ECs, stromal cells, and cancer cells via quantitative flow cytometry (qFlow) [5–15]. In particular, VEGFRs and NRP1 have been quantified in various healthy cells in vitro, including human umbilical vein ECs (HUVECs), human dermal fibroblasts, human dermal microvascular ECs (HDMECs), human dermal lymphatic microvascular endothelial cells, circulating ECs in human blood, and mouse fibroblasts [5–9, 12, 15]. Tumor VEGFR concentrations have been established in glioblastoma (patient-derived xenografts), breast cancer cells (in vitro and ex vivo in subcutaneous MDA-MB-231 tumors in nude mice), and ovarian cancer cells (in vitro) [6, 11, 13–15]. Additionally, VEGFRs have been measured on mouse hindlimb skeletal muscles in both healthy and ischemic conditions, ex vivo [6, 15, 16]. In these tissues and conditions, the qFlow approach provides surface biomarker data on cell heterogeneity while advancing statistical and computational biomarker analysis and offering insights into vascular and tumor biology [15, 17].

Despite these insights, receptors are not static on the plasma membrane: they exhibit dynamic trafficking between the plasma membrane and intracellular compartments [18]. Additionally, intracellular VEGFRs can bind to intracrine ligands [19], and endocytosed receptors can unbind and rebind, enabling intracellular signaling within cells [20]. Therefore, comprehensive whole-cell quantification of RTKs in ECs from different organs is an important yet underexplored area. Recently, techniques for whole-cell protein quantification were developed and applied to measure oxytocin receptors in human myometrial smooth muscle cells [21, 22]. However, this approach has not been applied to measuring RTKs.

To advance plasma membrane and whole-cell RTK (VEGFR1, VEGFR2, VEGFR3, NRP1, Axl, PDGFRα and β) measurement across organs, we used qFlow on ECs from different tissues: HDMECs (dermis), HUVECs (umbilical vein), HGMECs (kidney), HLiSMECs (liver), and HBMECs (brain), in vitro. We report the receptor concentrations both as ensembled averages over cell populations and on a cell-by-cell level to assess variation within those populations. Together, these measurements provide a foundational understanding of receptor compartmentalization across organ-specific ECs in vitro.

## Materials and Methods

### Cell culture

Primary kidney glomerular microvascular endothelial cells (HGMECs) (Cell Systems, ACBRI 127), primary liver sinusoidal microvascular endothelial cells (HLiSMECs) (Cell Systems, ACBRI 566), and primary human brain microvascular endothelial cells (HBMECs) (Cell Systems, ACBRI 376) were cultured in Complete Classic Medium (4Z0-500) supplemented with CultureBoost and Bac-Off antibiotic (Cell Systems, 4Z0-644). Primary human dermal microvascular endothelial cells (HDMECs) (ATCC, CRL-4060) underwent *PDGFRA* and *PDGFRB* double knock out (DKO) (to eliminate PDGFRα and β expression, respectively) via CRISPR/Cas9 at the Genome Engineering & Stem Cell Center at Washington University in St. Louis, MO. The efficiency of DKO was validated by next-generation sequencing (NGS) analysis (Table S1). The purpose of this gene modification is to validate the threshold of specific binding of qFlow. HDMECs (dermis; *PDGFRA^−/−^* and *PDGFRB^−/−^*) were cultured in Vascular Cell Basal Medium (ATCC, PCS100030) supplemented with microvascular endothelial cell growth kit -VEGF (i.e., with VEGF; ATCC, PCS110041) and penicillin streptomycin (10,000 U/mL) (Thermo Fisher Scientific, 15-140-122). Primary human umbilical vein endothelial cells (HUVECs; ATCC, PCS100013) were cultured in EGM-2 (Lonza, CC-3162) containing Bac-Off antibiotic (Cell Systems, 4Z0-644). Cells in passages 3–6 were used and maintained in a humidified incubator at 37 °C and with 5% CO_2_.

### Quantitative Flow Cytometry

ECs were seeded in T-75 flasks and grown to approximately 80% confluence. The cells were washed with Ca^2+^/Mg^2+^-free PBS, harvested in Corning CellStripper solution, a nonenzymatic cell-dissociation solution, and incubated for 15 min at 37 °C and 5% CO_2_. The disassociation was stopped by adding Ca^2+^/Mg^2+^-free PBS, and the cell suspension was centrifuged twice at 400×*g* for 5 min at 4 °C. The cells were resuspended at 2–4 ×10^6^ cells/mL in ice-cold stain buffer (PBS, bovine serum albumin, sodium azide). We added 25-µL aliquots of a single-cell suspension for plasma membrane staining and 50 μL for whole-cell staining to 96-well deep-well plates. The appropriate amount of human VEGFR1 phycoerythrin (PE)-conjugated antibody (R&D Systems, FAB321P), VEGFR2 PE-conjugated antibody (BioLegend, 359904), VEGFR3 PE-conjugated antibody (R&D Systems, FAB3492P), NRP1 PE-conjugated antibody (R&D Systems, FAB3870P), Axl PE-conjugated antibody (R&D Systems, FAB154P), PDGFRα PE-conjugated antibody (R&D Systems, FAB1264P), or PDGFRβ PE-conjugated antibody (R&D Systems, FAB1263P) was added to each sample well except the fluorescence-minus-one (FMO) control. The saturating concentrations of PE-conjugated antibodies to detect receptors on the plasma membrane were previously determined at 14 μg/mL for VEGFR1, 2, and 3; 7.1 μg/mL for NRP1; and 9.4 μg/mL for PDGFRα and PDGFRβ [5, 8]. The saturating concentration of Axl antibody was determined at the beginning of the study to be 4 ug/mL to detect plasma membrane receptors (Fig. 3C). For whole-cell receptor detection, the saturating concentration of VEGFR1 and VEGFR2 antibodies was 20 µg/mL for both (Fig. 3A, 3B), and PE-VEGFR3 antibody saturated at 30 µg/mL (Fig. 3C). For PDGFR antibodies the same concentration was applied as that of VEGFR antibodies. Samples were incubated in dark for 40 min at 4 °C and washed twice with stain buffer by centrifugation at 400×*g* for 5 min at 4 °C. Samples for plasma membrane receptor measurement were washed and resuspended in 100 µL stain buffer before analysis.

Samples for whole-cell receptor measurement were fixed using 2% paraformaldehyde for 20 min in the dark, washed, and permeabilized (0.5% Tween-20) for 20 min in the dark. Samples were washed twice with stain buffer containing 0.1% Tween-20. PE-conjugated antibodies were added to the samples for staining for 40 min in the dark, followed by three washes with stain buffer containing 0.1% Tween-20. Washed samples for whole-cell receptor measurement were resuspended in 50 µL stain buffer. The readouts were acquired by a Cytoflex S flow cytometer (Beckman Coulter, IN) and CytExpert software (Beckman Coulter, IN). Kaluza software (Beckman Coulter, IN) was used for data analysis. Quantibrite PE beads (BD Biosciences, 340495) were collected and analyzed under the same compensation and voltage settings as cell fluorescence data. Quantibrite PE beads are a mixture of beads with four different concentrations of conjugated PE molecules: low (474 PE molecules/bead), medium-low (5359 PE molecules/bead), medium-high (23,843 PE molecules/bead), and high (62,336 PE molecules/bead). A calibration curve that translated PE geometric mean to the number of bound molecules was determined using linear regression, *y* = *mx* + *c*, where y is log_10_ fluorescence and x is log_10_ PE molecules per bead (Fig. 1D). Receptor levels were calculated as described previously [7, 8, 10, 11].

**Fig. 1.**
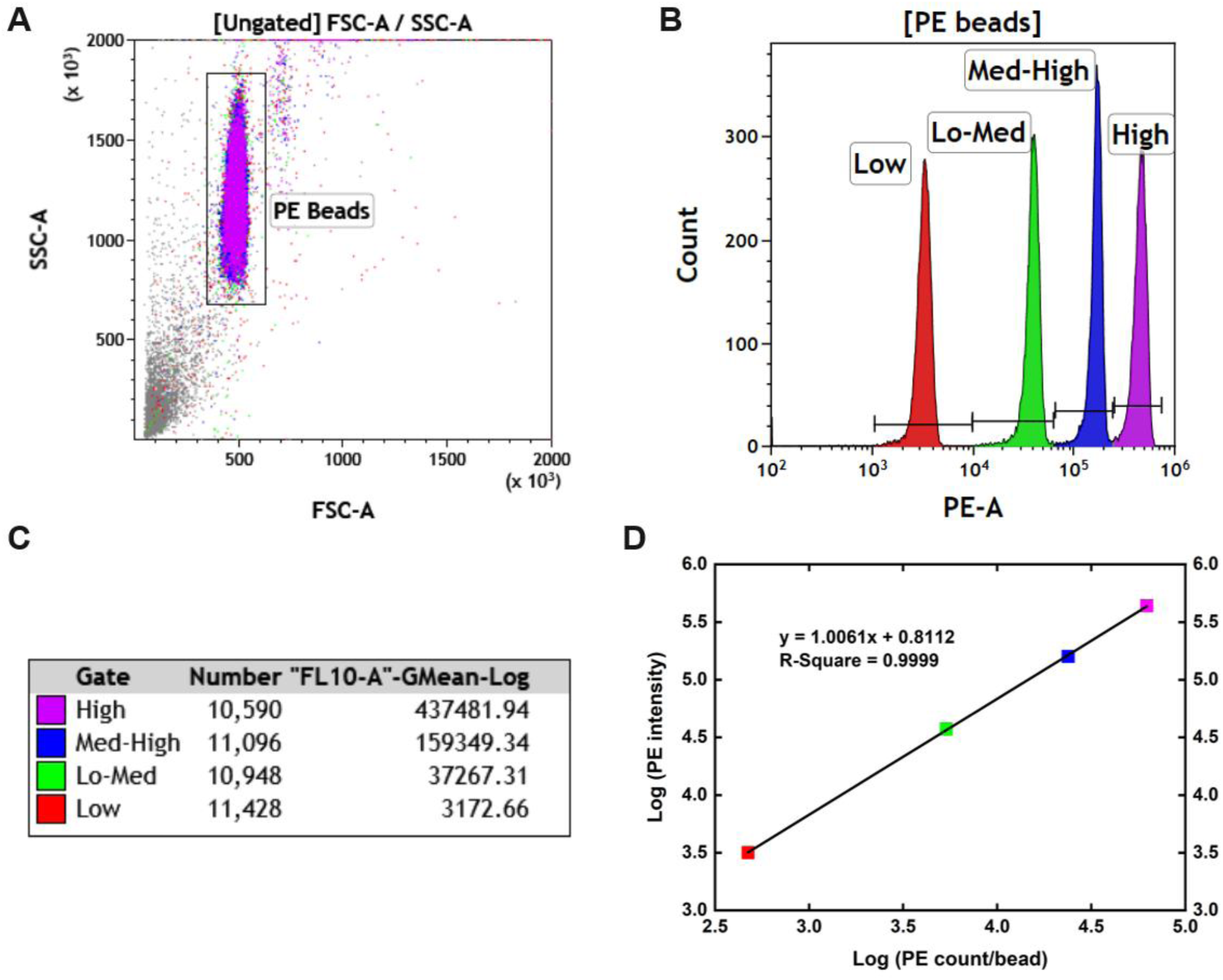
PE bead calibration curve based on the linear relationship between PE fluorescence and PE molecule intensities. (A) PE beads are gated on the FSC vs. SSC plot. (B) The four peaks are separately gated in the PE histogram, representing the four types of beads. (C) The histogram statistics show the geometric mean of PE intensity of each peak. (D) A linear PE bead calibration curve is generated by relating log_10_ PE molecules per bead to log_10_ fluorescence.

### Cell-by-Cell Analysis

Cell heterogeneity based on each examined receptor was analyzed from cell-by-cell PE fluorescence intensity as previously described [8, 9, 14, 16]. Briefly, a 2D histogram was generated to show the distribution of receptor numbers converted from EC fluorescence intensity using the PE bead calibration (Fig. 1D). A total of 5000 cells were randomly pooled from each cell population. Background noise was subtracted on the basis of the signal differences between labeled and unlabeled cells [5, 9]. The median and coefficient of variation (CV) are reported in Table 2. A two-sample Kolmogorov–Smirnov (K–S) test was performed in Matlab to determine if the distribution for each receptor and EC type was from a common distribution. In each case, the K–S test found the distributions to be significantly different (Table 2). To quantify cell-to-cell variation in RTK expression within each EC type, we used quadratic entropy (QE), which measures diversity by evaluating the weighted difference between expression levels across cells. The approach divides each cell-by-cell distribution into 500 equally spaced bins and calculates a sum of weighted differences between every two bins based on their average expression levels and distance [9, 23–25].

### Statistics

All data were acquired in at least three independent experiments. Ensemble averaged data are represented by mean ± standard error of the mean (SEM). One-way analysis of variance (ANOVA), multiple comparison post-hoc Tukey’s test, and power analysis of pair-comparison were performed.

## Results

### Whole-cell quantification to measure plasma membrane and intracellular RTKs

Prior qFlow studies measured plasma membrane RTKs [5, 7–9, 11]. We extended the traditional plasma membrane-focused qFlow studies to assess RTKs across the entire cell, including both plasma membrane and intracellular compartments using the whole-cell qFlow approach [21, 22] (Fig. 2). This approach involved staining intracellular proteins following fixation and permeabilization. Prior saturation studies helped determine appropriate antibody concentration ranges for plasma membrane labeling for VEGFRs, NRP1, and PDGRs [5, 8]. Here, we determined a saturating concentration of 4 μg/mL for plasma membrane Axl detection (Fig. 3C). For whole-cell quantification, we identified a saturating concentration of 20 μg/mL for both VEGFR1 (Fig. 3A) and VEGFR2 (Fig. 3B); as such, this saturation concentration of antibody is sufficient to fully detect VEGFRs.

**Fig. 2.**
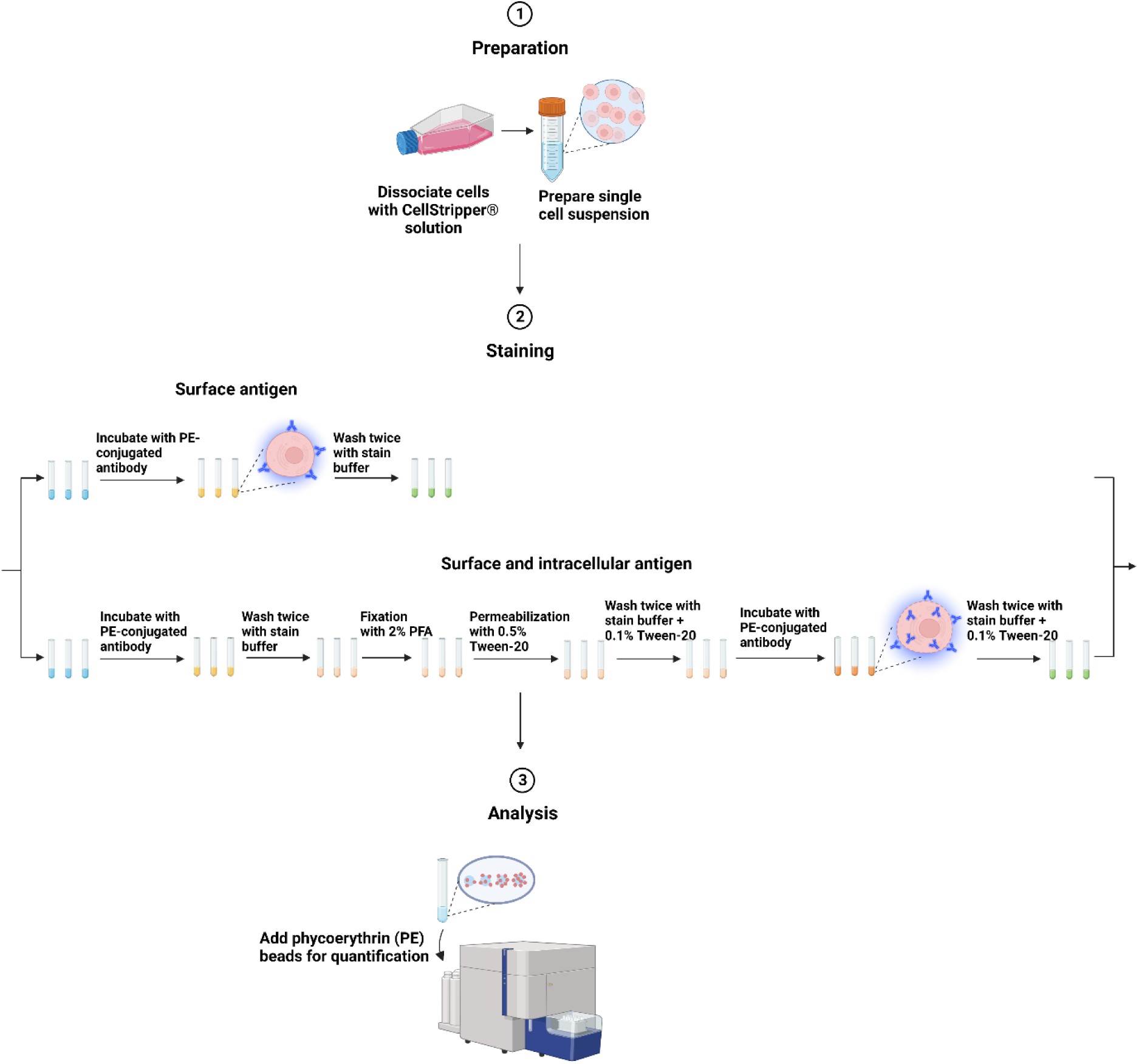
Schematic figure showing the workflow of quantitative flow cytometry measuring both surface and intracellular antigens. (Created with BioRender.com)

**Fig. 3.**
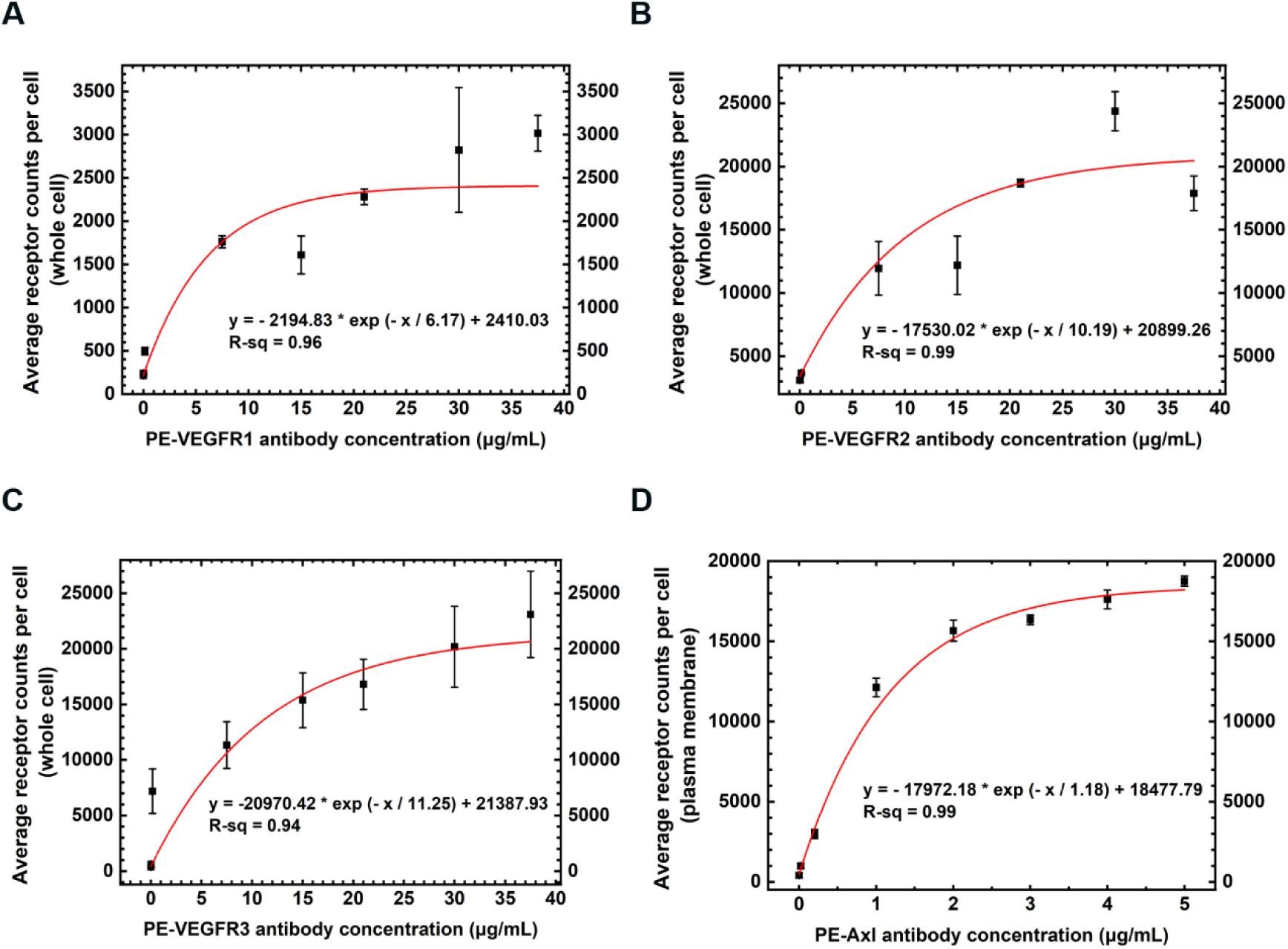
Saturation curves for PE-VEGFR1, PE-VEGFR2, and PE-VEGFR3 antibodies on the whole cell and PE-Axl antibody on the plasma membrane of ECs. (A) PE-VEGFR1 and (B) PE-VEGFR2 antibodies both saturated at 20 μg/mL on the whole cell of HUVECs with respect to an exponential fitting of normalized receptor density with increasing labeling concentration. (C) PE-VEGFR3 antibodies saturated at 30 μg/mL on the whole cell of HBMECs. Fitting equations were 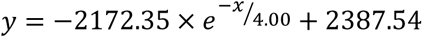, 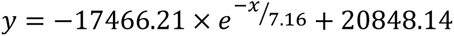, and 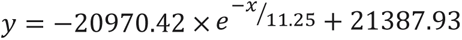, respectively. (D) PE-Axl saturated at 4 μg/mL on the plasma membrane of HBMECs with respect to an exponential fitting of normalized receptor density with increasing labeling concentration. The fitting equation was 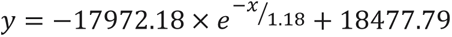.

### ECs exhibit organ-specific RTK concentrations at the plasma membrane and whole-cell levels

Using qFlow, we quantified RTK concentrations (VEGFR1, VEGFR2, VEGFR3, NRP1, Axl, and PDGFRα and β) on both the plasma membrane and whole cell across various EC types: HDMECs (dermis), HUVECs (umbilical vein), HGMECs (kidney), HLiSMECs (liver), and HBMECs (brain), respectively (Fig. 4; Table 1). Please note: HDMECs (dermis; *PDGFRA^−/−^* and *PDGFRB^−/−^*) were genetically modified to knock out both PDGFRα and β expression (see Methods). Our ensemble-averaged data show that the five EC types exhibited concentrations of each of the following receptors at the same order of magnitude: VEGFR1 (300–900 VEGFR1/cell on the plasma membrane and 2500–7500 VEGFR1/whole cell), VEGFR2 (4800–7500 VEGFR2/cell on the plasma membrane and 14,100–22,500 VEGFR2/whole cell), VEGFR3 (1200–4400 VEGFR3/cell on the plasma membrane and 32,400–62,000 VEGFR3/whole cell), NRP1 (26,000–39,700 NRP1/cell on the plasma membrane), and Axl (6900–12,200 Axl/cell on the plasma membrane). Notably, both PDGFRα an PDGFRβ were minimally expressed, across all four cell types possessing intact PDGFRA and PDGFRB genes; and the DKO HDMECs expressed few to no molecules on the plasma membrane (< 400 PDGFR/cell) and low levels across the whole cell (< 4000 PDGFR/cell) (Fig. S1). The 400 receptors/cell threshold for antibody-receptor binding was established by measuring nonspecific binding via an across-species antibody reactivity study, hPDGFR antibody binding vs mPDGFR antibody binding on mouse fibroblasts [7]. During whole-cell staining, nonspecific binding increased the total PDGFR concentrations quantified by about tenfold compared to plasma membrane levels (Fig. S1). However, the PDGFR double knockout gene modification, does not affect the expression or localization of other RTKs, as confirmed by comparable receptor concentrations observed between DKO HDMECs and wild-type HDMECs [8].

**Fig. 4.**
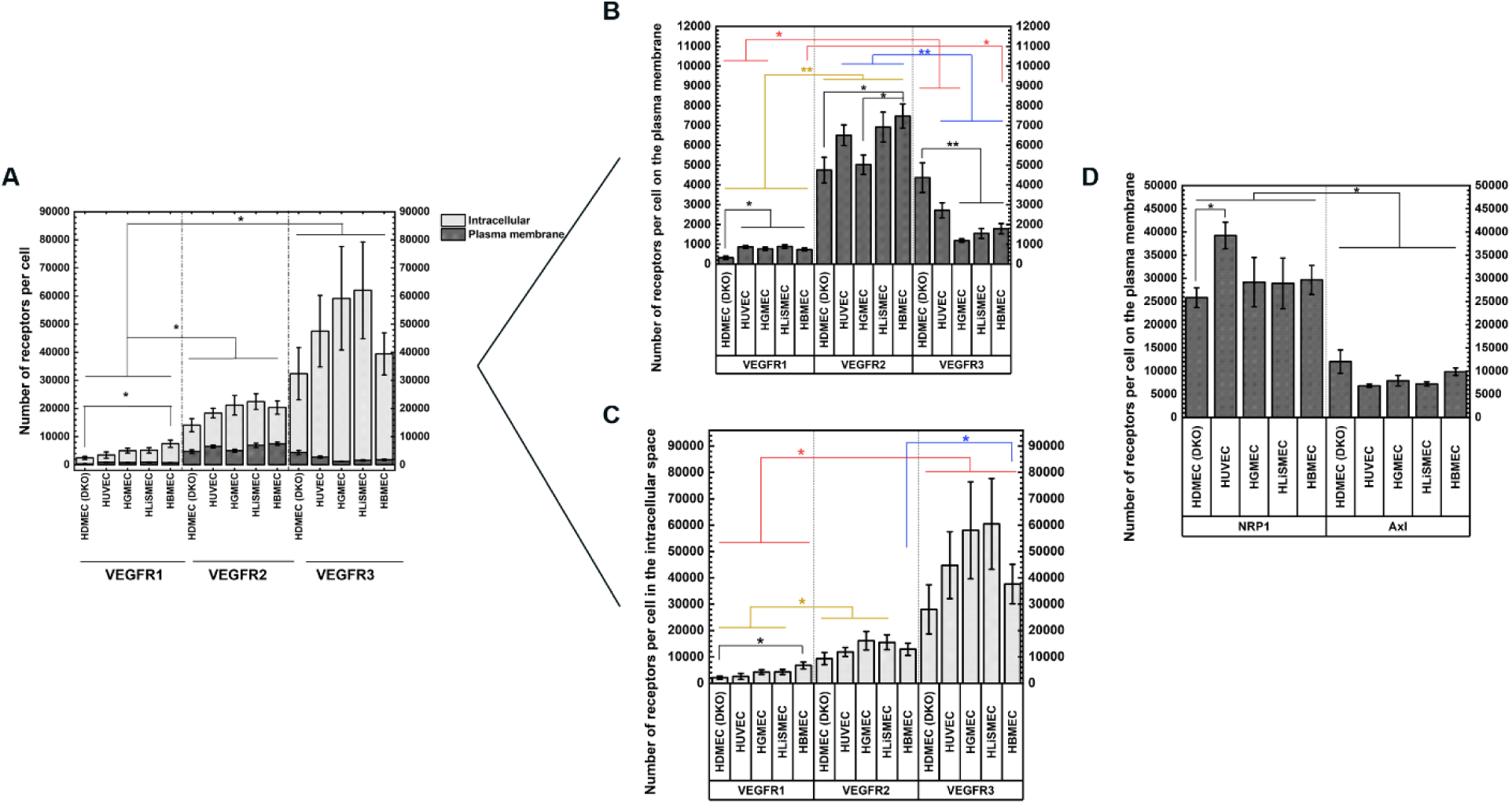
Plasma membrane and intracellular distributions of RTKs on different endothelial cells (ensemble average ± SEM). (A–C) VEGFR1, VEGFR2, and VEGFR3; (D) NRP1 and Axl. HDMEC (DKO), primary human dermal microvascular endothelial cells with *PDGFRA* and *PDGFRB* double knockout; HUVECs, primary human umbilical vein endothelial cells; HGMECs, primary kidney glomerular microvascular endothelial cells; HLiSMECs, primary liver sinusoidal microvascular endothelial cells; HBMECs, primary human brain microvascular endothelial cells.

**Table 1.** Number of each RTK on endothelial cells of various origins.

| Cell type | Receptor | Number/cell on the plasma membrane (Mean ± SEM) | Number/cell, whole cell (Mean ± SEM) | Percentage on the plasma membrane |
| --- | --- | --- | --- | --- |
| HDMEC (dermis; *PDGFRA*<sup>-/-</sup> and *PDGFRB*<sup>-/-</sup>) | VEGFR1 | 300 ± 90 | 2500 ± 600 | 12% |
| | VEGFR2 | 4800 ± 600 | 14,100 ± 1800 | 34% |
| | VEGFR3 | 4400 ± 700 | 32,400 ± 9800 | 14% |
| | NRP1* | 26,000 ± 1100 | - | - |
| | Axl* | 12,200 ± 1300 | - | - |
| HUVEC (umbilical vein) | VEGFR1 | 900 ± 70 | 3500 ± 1100 | 26% |
| | VEGFR2 | 6500 ± 500 | 18,400 ± 2100 | 35% |
| | VEGFR3 | 2700 ± 400 | 47,500 ± 12,600 | 6% |
| | NRP1* | 39,700 ± 2900 | - | - |
| | Axl* | 6900 ± 400 | - | - |
| HGMEC (kidney) | VEGFR1 | 800 ± 80 | 5100 ± 800 | 16% |
| | VEGFR2 | 5000 ± 500 | 21,200 ± 3600 | 24% |
| | VEGFR3 | 1200 ± 90 | 59,200 ± 18,300 | 2% |
| | NRP1* | 29,300 ± 5200 | - | - |
| | Axl* | 7900 ± 1000 | - | - |
| HLISMEC (liver) | VEGFR1 | 900 ± 90 | 5100 ± 900 | 18% |
| | VEGFR2 | 6900 ± 800 | 22,500 ± 2600 | 31% |
| | VEGFR3 | 1800 ± 300 | 62,000 ± 17,000 | 3% |
| | NRP1* | 29,000 ± 5300 | - | - |
| | Axl* | 7200 ± 400 | - | - |
| HBMEC (brain) | VEGFR1 | 700 ± 70 | 7500 ± 1300 | 9% |
| | VEGFR2 | 7500 ± 600 | 20,400 ± 1900 | 37% |
| | VEGFR3 | 1800 ± 300 | 39,400 ± 7200 | 5% |
| | NRP1* | 29,900 ± 2900 | - | - |
| | Axl* | 9900 ± 900 | - | - |

\*Whole-cell NRP1 and Axl levels are not shown because we found NRPs and Axl to be sensitive to cell fixation and permeabilization processes during whole-cell staining.

Of the RTKs measured, VEGFR1 exhibited the lowest concentrations on the plasma membrane, < 1,000 VEGFR1/cell in all five cell types investigated. HDMECs (dermis; *PDGFRA^−/−^* and *PDGFRB^−/−^*) had significantly lower plasma membrane VEGFR1 levels, at 60% of the average concentration compared to the other four cell types: HBMECs (brain), HGMECs (kidney), HLiSMECs (liver), and HUVECs (umbilical vein) (p < 0.05; all p-values were calculated by one-way ANOVA with post hoc Tukey’s test) (Fig. 4B; Table 1).

Among the cell types studied, there were differences in the percentage of VEGFR1 found on the plasma membrane versus the entire cell. HBMECs (brain) showed the lowest percentage of total VEGFR1 located on the plasma membrane, at 9% of the total VEGFR1 present. In contrast, HUVECs (umbilical vein) showed the highest proportion, with 26% of the total VEGFR1 localized to the plasma membrane. Following HUVECs, the HLiSMECs (liver) showed 18%, HGMECs (kidney) had 16%, and HDMECs (dermis; *PDGFRA^−/−^*and *PDGFRB^−/−^*) exhibited 12% localization to the plasma membrane (Table 1). In summary, the order of cell types by the percentage of VEGFR1 located on the plasma membrane was, from highest to lowest, HUVECs > HLiSMECs, HGMECs, HDMECs > HBMECs.

At the whole-cell level, among the VEGFRs, VEGFR1 concentrations were the lowest: < 8000 molecules/cell across all cell types investigated. As in the plasma membrane analysis, HDMECs (dermis; *PDGFRA^−/−^* and *PDGFRB^−/−^*), compared with the other four cell types, exhibited the lowest whole-cell concentration of VEGFR1. Notably, the whole-cell concentration of VEGFR1 in HDMECs (dermis; *PDGFRA^−/−^* and *PDGFRB^−/−^*) was 67% lower than that observed in HBMECs (brain) (p < 0.05; Fig. 4A). The remaining cell types, HLiSMECs (liver), HGMECs (kidney), and HUVECs (umbilical vein), demonstrated similar whole-cell concentrations of VEGFR1 protein, averaging 4600 VEGFR1/cell (Table 1).

VEGFR2 demonstrated consistent localization on the plasma membrane across the five cell types tested. VEGFR2 demonstrated significantly higher plasma membrane localization compared to VEGFR1 (p < 0.01); indeed, at least one order of magnitude higher (Fig. 4B; Table 1). Among all the receptors studied, VEGFR2 exhibited the highest percentage of localization on the plasma membrane, with an average of 32% of total VEGFR2 being located on the plasma membrane. In ranked order, HDMECs (dermis; *PDGFRA^−/−^* and *PDGFRB^−/−^*) and HGMECs (kidney) had the lowest concentrations of VEGFR2 on the plasma membrane (4800–5000 molecules, p < 0.05 for both), while HBMECs (brain) had the highest VEGFR2 concentration on the plasma membrane, with 7500 ± 600 molecules (p < 0.05; Fig. 4B, Table 1).

Whole-cell VEGFR2 protein expression was consistent across all investigated cell types, with concentrations ranging from 14,100–22,500 VEGFR2/cell (Table 1). This is approximately one order of magnitude higher than that of VEGFR1 at the whole-cell level (p < 0.05). Notably, although HDMECs (dermis; *PDGFRA^−/−^* and *PDGFRB^−/−^*) exhibited the lowest VEGFR2 concentrations among the cell types, the observed differences did not reach statistical significance (0.06 ≤ p ≤ 0.9; Fig. 4A).

The VEGFR3 plasma membrane concentrations were intermediate, relative to VEGFR1 and VEGFR2, but VEGFR3 showed the lowest percentage of plasma membrane localization of all receptors studied. Plasma membrane VEGFR3 levels in HDMECs (dermis; *PDGFRA^−/−^* and *PDGFRB^−/−^*), HUVECs (umbilical vein), HGMECs (kidney), and HBMECs (brain) were significantly higher than those of VEGFR1 in the same cell types (p < 0.01). Whereas plasma membrane VEGFR3 concentrations in HGMECs (kidney), HLiSMECs (liver), and HBMECs (brain) were significantly lower than plasma membrane VEGFR2 concentrations (p < 0.01). HDMECs (dermis; *PDGFRA^−/−^* and *PDGFRB^−/−^*) had the highest plasma membrane concentration of VEGFR3 (p < 0.01), with 4400 ± 700/cell, and the highest percentage of total VEGFR3 on the plasma membrane, at 14% (Table 1), displaying a 1.6-fold higher plasma membrane concentration of VEGFR3 than that observed in HUVECs (umbilical vein) (Fig. 4B; Table 1). Furthermore, HUVECs (umbilical vein) had a VEGFR3 concentration 1.7-fold higher than that observed in HGMECs (kidney), HLiSMECs (liver), and HBMECs (brain) (p < 0.01; Fig. 4B).

Whole-cell VEGFR3 concentrations were universally high, ranging from 32,400–62,000 VEGFR3/cell. Despite this wide whole-cell VEGFR3 concentration range, no significant differences were observed across the five cell types (0.22 ≤ p ≤ 0.9) (Fig. 4A). However, it is important to note that this whole-cell VEGFR3 concentration range was significantly higher than that of whole-cell VEGFR1 by approximately 10-fold (p < 0.05; Fig. 4A, Table 1). While whole-cell VEGFR3 levels were higher than any other whole-cell RTK levels assayed, in most cell types 94–98% was intracellular, and thus only a small fraction of the total was located on the plasma membrane: 6% in HUVECs (umbilical vein), 5% in HBMECs (brain), 3% in HLiSMECs (liver), and 2% in HGMECs (kidney) (Table 1).

Plasma membrane NRP1 was significantly more abundant than any VEGFR, with concentrations exceeding 26,000 per cell in all cell types (p <0.05). HUVECs displayed the highest NRP1 concentration (39,700 ± 2900/cell on the plasma membrane), which was significantly higher than the level observed on the plasma membrane of HDMECs (dermis; *PDGFRA^−/−^* and *PDGFRB^−/−^*), representing a 1.5-fold difference (p < 0.05; Fig. 4D). No significant difference was observed across plasma membrane NRP1 levels on HUVECs (umbilical vein), HGMECs (kidney), HLiSMECs (liver), and HBMECs (brain) (Fig. 4D). Axl was detected at lower concentrations than NRP1 on the plasma membrane, ranging from 6900–12,200/cell (p < 0.05, Table 1) It is important to note that the whole-cell protein expression levels of NRP1 and Axl are not presented in this study. This is further explained in the discussion.

Cell-by-cell analysis indicated that the receptor concentrations followed log-normal distributions. This pattern, observed for both plasma membrane and whole-cell receptor levels, exhibited positive skewness — indicating that most cells had lower receptor numbers with a minority possessing much higher concentrations. This skewness suggests that a few cells may play a disproportionately larger role in receptor-mediated processes. Additionally, positive kurtosis was noted, indicating a steeper peak and heavier tail than a normal distribution, further emphasizing the variability within cell populations (Table 2; Fig. 5) [26]. Despite these shared characteristics of skewness and kurtosis, peaks within these distributions varied in position and spread across different EC types.

**Table 2.**
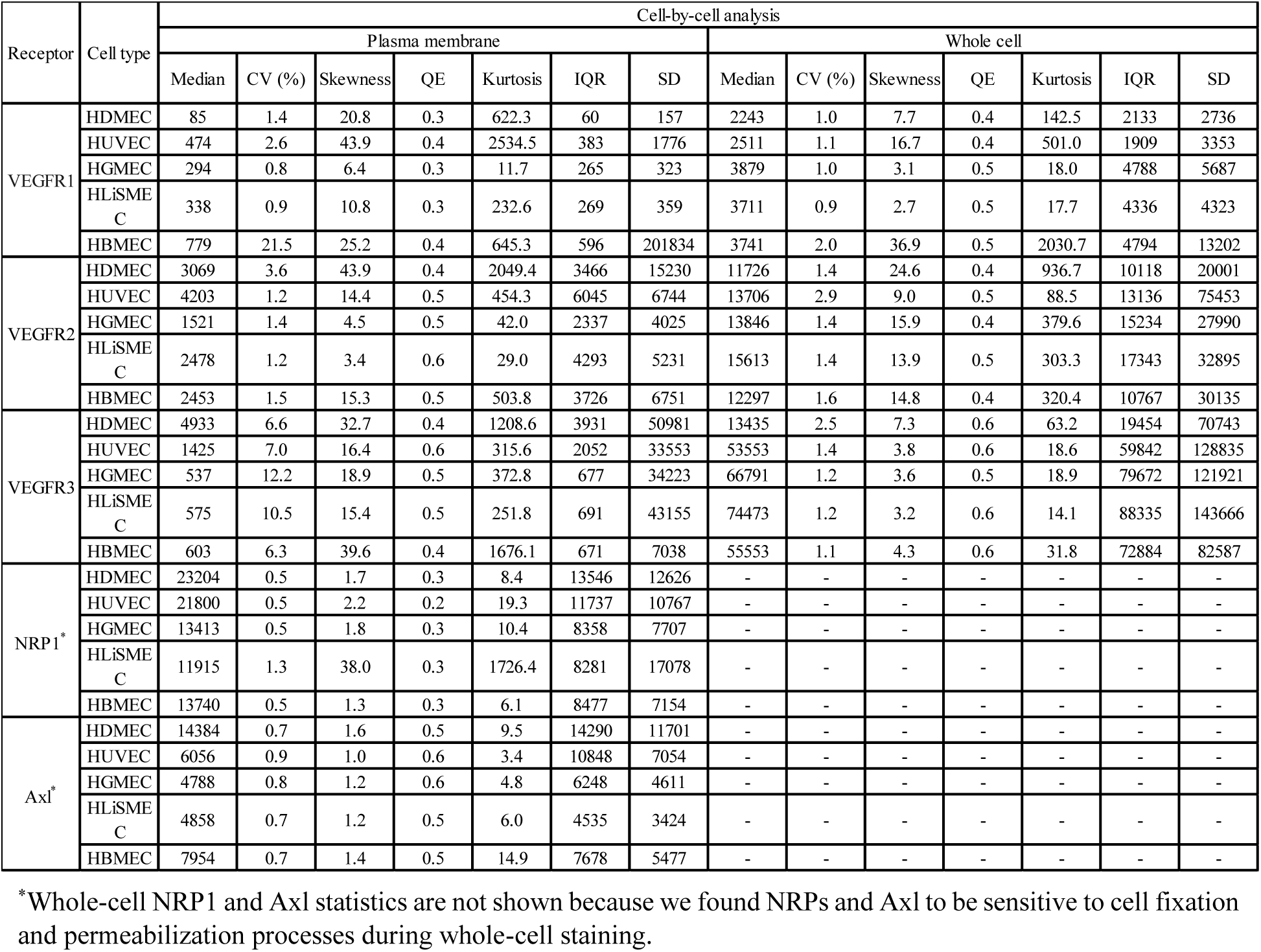
Receptor statistics of cell-by-cell analysis.

**Fig. 5.**
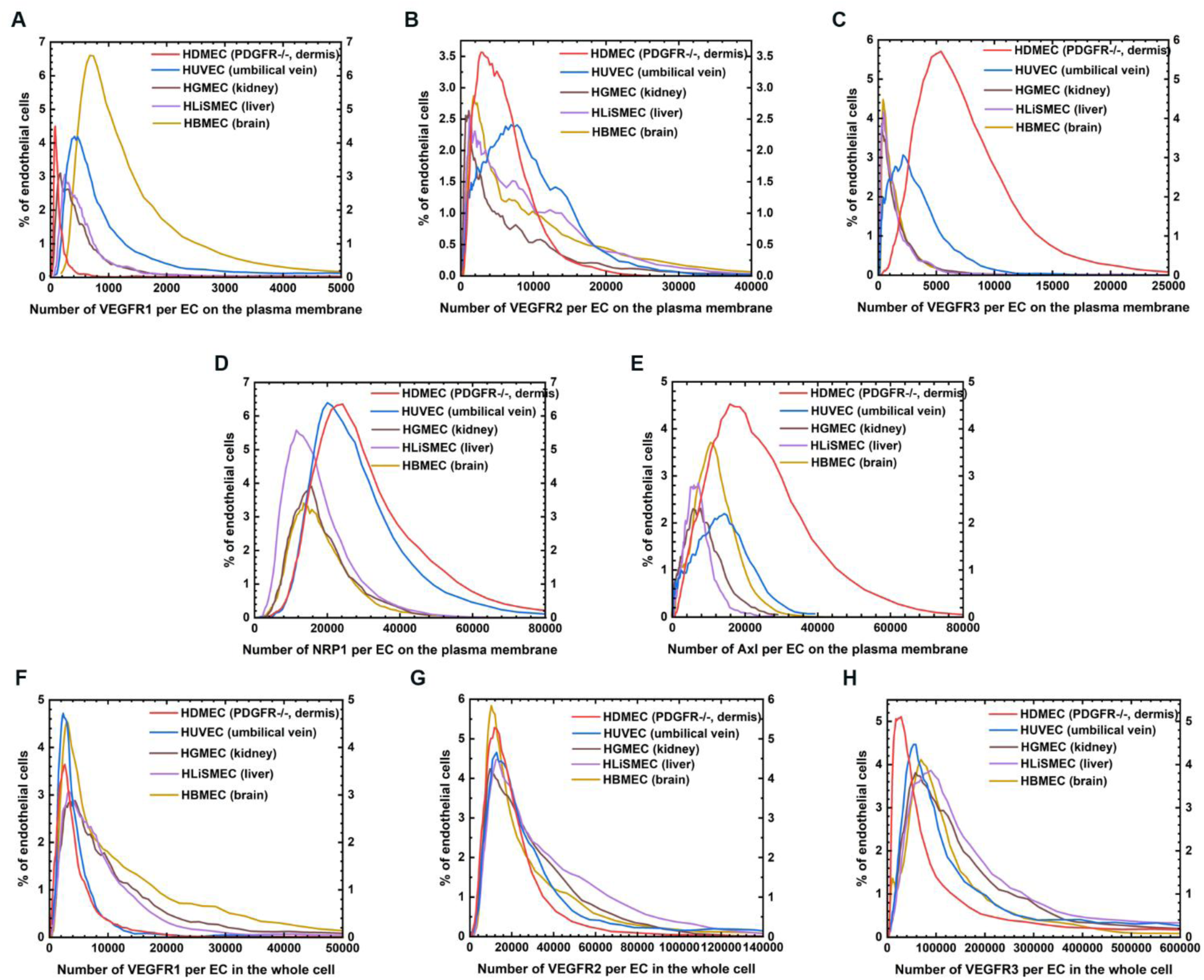
Cell-by-cell analysis of receptor distributions on the plasma membrane (A–E) or across the whole cell (F–H) of ECs from the five organs. (A) VEGFR1, (B) VEGFR2, (C) VEGFR3, (D) NRP1, and (E) Axl on the plasma membrane. (F) VEGFR1, (G) VEGFR2, (H) VEGFR3 across the whole cell.

Among the tested cell types, HBMECs (brain) exhibited a particularly broad distribution in plasma membrane VEGFR1 concentrations. This was evidenced by the largest coefficient of variance (CV) and interquartile range (IQR), with a right-shifted peak (Fig. 5A; Table 2). This suggests a greater variability in VEGFR1 distribution, possibly reflecting the dynamic nature of VEGFR1 in brain microvascular endothelia where localized signaling might require rapid receptor turnover or varied availability[27].

In contrast, HDMECs (dermis; *PDGFRA^−/−^* and *PDGFRB^−/−^*) displayed the highest variability for plasma membrane VEGFR2, demonstrated by their high CV, skewness, kurtosis, and standard deviation (SD). Whereas on HUVECs (umbilical vein), the plasma membrane VEGFR2 distribution had a broader spread, with the highest IQR (Fig. 5B; Table 2).

Plasma membrane VEGFR3 exhibited distinct distribution peaks among different cell types (Fig. 5C). HDMECs (dermis; *PDGFRA^−/−^* and *PDGFRB^−/−^*) showed the broadest distribution (largest SD and IQR) with a right-shifted peak (highest median), indicating higher medians compared to other cell types like HUVECs (umbilical vein), which showed a median-aligned peak. HGMECs (kidney), HLiSMECs (liver), and HBMECs (brain) were clustered towards lower medians and IQRs (Table 2), suggesting a more uniform but moderate expression profile of plasma membrane VEGFR3 in these cells.

For plasma membrane NRP1, HDMECs (dermis; *PDGFRA^−/−^* and *PDGFRB^−/−^*) and HUVECs (umbilical vein) shared a similar range in CV, IQR, and SD), reflecting clustered peaks with similar medians, indicative of a relatively stable and uniform NRP1 distribution among these cell types (Fig. 5D; Table 2). In contrast, plasma membrane Axl levels in HDMECs (dermis; *PDGFRA^−/−^*and *PDGFRB^−/−^*) showed a broader range with the largest IQR and SD, and a right-shifted peak, reflecting a notably higher median (Fig. 5E).

The distributions displayed a consistent pattern of homogeneity within each cell type, exhibiting a tightly clustered distribution with a single peak for each receptor, and supported by QE values consistently below 0.7, indicating low heterogeneity in both the plasma membrane and whole-cell compartments (Table 2). QE < 0.7 is considered to indicate low heterogeneity on the basis of the variation in receptor expression, and QE > 0.7 high heterogeneity [8, 9, 14]. Notably, the whole-cell RTK peaks were more aligned than the plasma membrane peaks (Fig. 5F–5H), which was consistent with the absence of statistically significant differences between cell types at the whole-cell level for each receptor type (Fig. 4C). These findings collectively suggest a high degree of homogeneity among the cultured ECs, underscoring the consistent RTK protein expression patterns across different cell types.

## Discussion

Our study presents a comprehensive analysis of RTK protein abundance and distribution across ECs derived from various organs. We highlight three major findings: (1) Consistent RTK Levels Across Organ-Derived ECs: Despite the inherent phenotypic heterogeneity of ECs, we observed that the RTK expression levels on the plasma membrane and at the whole-cell level are of the same order of magnitude across different cell types for each RTK studied. This is indicative of a fundamental consistency in RTK protein abundance independent of organ origin. (2) Differential Localization Patterns: Our analysis uncovered distinct localization patterns among VEGFRs. Specifically, VEGFR1 and VEGFR3 were primarily intracellular, whereas VEGFR2 showed significant presence on the plasma membrane. This differential localization suggests unique functional roles mediated by receptor compartmentalization. (3) Minimal Variation in RTK Expression Corroborates EC Homogeneity in Culture: The RTK expression patterns exhibited low heterogeneity, as indicated by the low QE values. This suggests that in vitro conditions may mitigate the diverse expression profiles typically observed in vivo, emphasizing the importance of environmental context in EC phenotype.

### VEGFR1 Role beyond A Decoy Receptor

The traditionally held view of VEGFR1 as a “decoy” receptor is challenged by our findings and existing literature [9, 11, 12, 15, 16, 28], necessitating a reevaluation of its functional roles. Conventionally, VEGFR1 has been depicted as sequestering VEGF-A away from the more pro-angiogenic VEGFR2 due to its higher affinity for VEGF-A [28, 29]. This competitive binding model is supported by the fact that VEGFR1 possesses markedly lower kinase activity [30, 31]. Furthermore, knockout studies define VEGFR1 as inhibiting vascularization, where the absence of VEGFR1 results in disorganized and overgrown vessels [32, 33].

In our study, the number of VEGFR1 molecules on the plasma membrane was approximately 90% lower than that of VEGFR2, and the overall cellular VEGFR1 expression was about 60% lower than VEGFR2. These findings are in line with previous flow cytometry and biotin labeling data from in vitro studies, confirming the limited plasma membrane expression of VEGFR1 [5, 7, 34]. However, our observations suggest that classifying VEGFR1 merely as a decoy might overlook its potential active roles in angiogenesis.

Indeed, the substantial proportion of intracellular VEGFR1 may hint at unrecognized facts of this receptor, possibly engaging in intracrine signaling pathways or receptor recycling processes[35]. Such mechanisms might influence vascular responsiveness to angiogenic cues under certain physiological or pathological conditions. Computational models have suggested that membrane-associated VEGFR1 is vital for regulating free VEGF levels in the bloodstream, thereby influencing cell migration and stabilization of vascular networks [15]. Experimental data further supports a broader role for VEGFR1, where increased VEGFR1 expression has been linked with angiogenic responses. For instance, during reperfusion following hindlimb ischemia, a 2-fold increase in plasma membrane VEGFR1 was observed (2360 ± 220 molecules/cell), highlighting its possible involvement in vascular remodeling processes [16]. Moreover, a 10-fold elevation of plasma membrane VEGFR1 was observed in breast cancer ECs, with an average of 15,000 VEGFR1/cell observed on the plasma membrane after 3 weeks of tumor growth, further indicating that increased VEGFR1 levels are involved in active angiogenesis [11]. Similarly, human dermal fibroblasts displayed notable VEGFR1 expression (∼2700 VEGFR1/cell) on the plasma membrane when actively migrating in coculture with HUVECs in vitro, compared with no to low expression in monocultured fibroblasts [9].

Another notable point is that the low VEGFR1 plasma membrane expression observed in this study and in prior in vitro studies correspond to ECs not in active angiogenesis [5, 7, 34]. For example, in blood samples from menopausal and postmenopausal females, only low-VEGFR circulating ECs could be observed (approximately 3000 VEGFR1/cell); in contrast, in premenopausal females, where angiogenesis occurs monthly via menarche both low- and high-VEGFR (138,000 VEGFR1/cell) cell populations were observed [12]. Therefore, the low-plasma membrane VEGFR1 also prompts a shift in perspective, encouraging exploration into alternate, potentially intracrine signaling roles and its contribution to endothelial cell dynamics beyond the classical decoy paradigm.

### Stable VEGFR2 Protein Expression

Our study reveals that VEGFR2 levels remain remarkably stable across diverse conditions, suggesting robust internal regulatory mechanisms. Quantitatively, VEGFR2 presence was significantly higher than that of VEGFR1, both on the plasma membrane and in whole-cell measurements, with consistent levels between 4800–7500 molecules/cell on the plasma membrane and 14,100–22,500 molecules/cell throughout the whole cell. These findings align with previous in vitro research demonstrating stable VEGFR2 expression across various states, including healthy, tumor, and disease contexts [6, 14].

Further supporting this stability hypothesis is the fact that similar VEGFR2 protein expression levels on the plasma membrane have been observed in monolayer and spheroid-derived ovarian cancer cell lines [13]. In contrast, early-stage tumor ECs displayed dramatically elevated plasma membrane VEGFR1 levels (10-fold higher plasma membrane) compared to normal ECs, yet VEGFR2 levels remained constant with tumor growth, within the range of 1200–1700 molecules/cell [11]. Comparable VEGFR2 stability was also observed in actively angiogenic HUVECs cocultured with human dermal fibroblasts and ruing ex vivo ischemic conditions, where other receptor levels, like VEGFR1 significantly fluctuated [9, 16]. Furthermore, among all receptors analyzed, VEGFR2 exhibited minimal variation in cell-by-cell distribution (Fig. 5B, 5G), matching previous findings [9, 14], and reinforcing the notion of its tightly regulated expression.

The VEGFR2 stability could be attributed to its slower trafficking rates; involving synthesis, internalization, recycling, and degradation; compared with VEGFR1 [35]. The distribution of VEGFR2 protein is more balanced between the plasma membrane and the intracellular space. Moreover, VEGFR2 synthesis compensates for its degradation [35]. By contrast, the majority of VEGFR1 protein is located inside the ECs. When extracellular ligands are brought in and released from internalized ligated receptors, the numerous internal VEGFR1 molecules are activated and participate in protein dynamics and signaling [35].

Despite the consistent measurements of plasma membrane VEGFR2 numbers here and in previous work [6], the percentage of VEGFR2 located on the plasma membrane (35% in HUVECs) appears to be lower than what had been reported (50%–60% in HUVECs) using biotin labeling [34, 36]. This discrepancy underscores the importance of performing saturation analysis for quantitative work [5, 8] to accurately capture receptor numbers. The difference also suggests an opportunity to examine any sensitivity differences between qFlow and biotin labeling techniques.

### VEGFR3 Role in Vasculature

Traditionally recognized for its role in lymphatic vessel development, VEGFR3 levels have emerged in our study as significant within vascular ECs [37, 38]. Our data reveal that VEGFR3 is expressed more abundantly at the whole-cell level than both VEGFR1 and VEGFR2. However, intriguingly, it exhibits the least localized to the plasma membrane (2–14%), suggesting a predominantly intracellular role (Fig. 6; Table 1, S2).

**Fig. 6.**
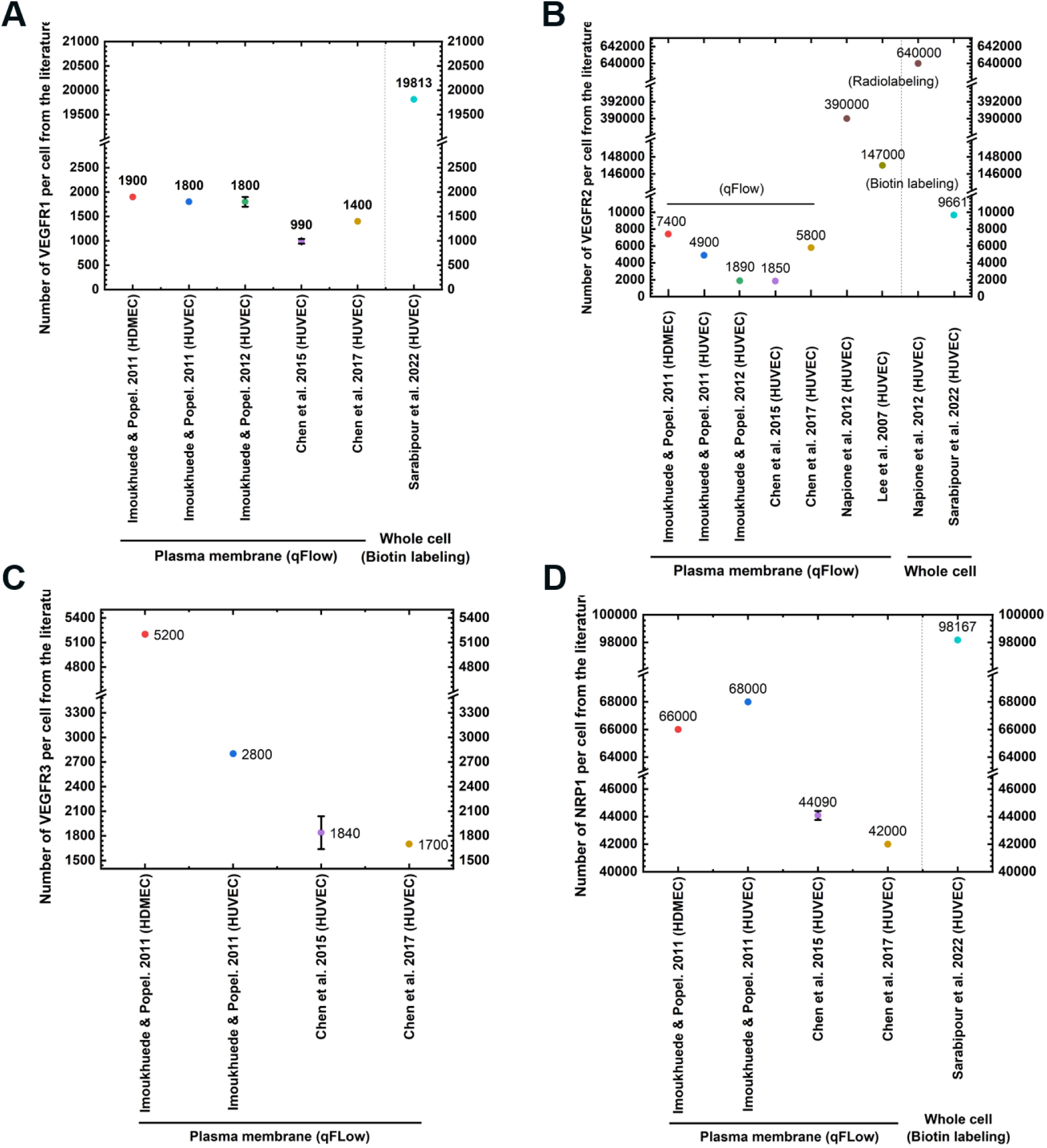
RTK quantification through qFlow, radio-, and biotin labeling, in vitro, from the literature. **(A)** VEGFR1, **(B)** VEGFR2, **(C)** VEGFR3, and **(D)** NRP1 have been quantified using qFlow, by radiolabeling, or by biotin labeling on the plasma membrane or at the whole-cell level [5–8, 34, 36, 87]. The numbers are summarized.

Within the vasculature, VEGFR3 impacts vascular permeability, as demonstrated by its role in modulating VEGFR2 activity and expression, particularly in the skin and kidneys [39]. We found that HDMECs (dermis; *PDGFRA^−/−^* and *PDGFRB^−/−^*) showed the highest concentration of VEGFR3 on the plasma membrane and the lowest concentration of plasma membrane VEGFR2. This suggests a VEGFR2– VEGFR3 balance potentially regulating vascular permeability. This observation invites further investigation into how these receptors interact to influence endothelial barrier functions.

Moreover, we also showed high whole-cell expression of VEGFR3 protein, approximately 60,000 molecules/cell, in HGMECs (kidney). This supports the established VEGFR3 role in modulating endothelial permeability and mediating glomerular microvasculature formation [39, 40]. Similarly, HLiSMECs (liver) had the highest whole-cell concentration of VEGFR3, consistent with the finding that VEGFR3 plays a critical role in liver vascular development [41]. The VEGF-C/VEGFR3 signaling axis is known to facilitate liver sinusoidal growth, inducing VE-cadherin endocytosis via SRC-mediated phosphorylation and upregulating VEGFR2 activation. Loss of VEGFR3 signaling causes reduced fetal liver size [41]. Our study thus provides quantitative VEGFR3 data that, when coupled with established vascular knowledge, sheds light on VEGFR3’s contributions: broadening its known role in lymphatic development and highlighting its potential for intracellular targeting to modulate vascular development and permeability across tissues.

### High NRP1 Expression and Its Implications for Angiogenesis

Our study found that NRP1 is highly expressed on the plasma membranes of ECs, with notably higher concentrations than VEGFR2 by 4–6 fold. Specifically, HUVECs displayed the highest NRP1 levels at approximately 39,700 ± 2900 molecules per cell (Table 1), aligning with prior observations and emphasizing the predominance of NRP1 in venous ECs (Fig. 6; Table S2) [5, 7, 42].

This elevated NRP1 concentration suggests its significant role in angiogenesis, primarily through its capacity to sequester VEGF-A, thus modulating the availability of this key growth factor [43, 44]. NRP1 achieves this function via post-translational modifications with glycosaminoglycan (GAG) chains, which facilitate the clustering of NRP1 and enhance VEGF-A binding, effectively competing with VEGFR2 [26, 43]. This sequestration mechanism is pivotal, as it regulates VEGF-A activity and mitigates abnormal angiogenesis, marking NRP1 as a compelling target for therapeutic intervention [26, 44].

Moreover, the function of NRP1 extends beyond angiogenesis. Its roles in maintaining endothelial homeostasis and preventing inflammation are well-documented, with studies showing that NRP1 loss leads to TGF-β pathway activation and junctional instability through VE-cadherin disruption [45, 46]. These functions are reinforced by our findings, suggesting that NRP1’s abundant expression is essential not only for vascular growth modulation but also for preserving vascular integrity and responsiveness.

### Axl: A Consistent Target for Anti-Angiogenic Therapy in Cancer

Our study identified Axl as being abundantly expressed on the plasma membranes of ECs, with concentrations ranging from 6900 to 12,200 molecules per cell — levels comparable to VEGFR2. This massive expression is particularly pronounced in HDMECs (dermis; *PDGFRA^−/−^* and *PDGFRB^−/−^*). The high level of Axl expression reflects its potential as a promising candidate for evaluating Axl-targeted therapies in angiogenesis-related pathologies [13], especially given its role in facilitating the migration and invasion of cancer cells, such as those seen in melanoma [47].

Our findings highlight the contrast between more homogeneous Axl expression in these studied ECs and the heterogeneity observed in ovarian cancer cells, where Axl levels varied significantly across different lines and culture conditions—from 350 to 33,000 molecules per cell [13]. This variation is reflective of the diverse genomic backgrounds of cancer patients and underscores the need for tailored therapeutic approaches to address the unique proteomic landscapes seen in tumors.

### Whole-Cell NRP1 and Axl Data Are Excluded because of The Sensitivity of These Proteins to Staining Processes

We have not reported whole-cell protein expression levels of NRP1 and Axl, because these proteins are sensitive to cell fixation and permeabilization processes during whole-cell staining. The paraformaldehyde used in fixation can lower the binding affinities of antibodies to receptors and cause morphological changes to cells and receptors that hinder antibody access to the receptors [48]. As a result, accurate whole-cell quantitation of NRP1 and Axl protein expression was not feasible using the established staining methods. Future studies should examine alternate whole-cell receptor quantification approaches, such as biotin or radio labeling.

### PDGFR Expression Dynamics in ECs Hint at The Impacts of The Microenvironment on Protein Levels

PDGFRs have not been found to be expressed by ECs in monoculture in vitro (Fig. S1). The numbers of PDGFRs measured on the five cell types were below the qFlow detection threshold for significant antibody– receptor binding (< 400 PDGFR/cell on the plasma membrane and < 4000 PDGFR/cell across the whole cell) (Fig. S1). However, studies have shown significant expression in ECs when cocultured with fibroblasts or monocultured on a surface coated with fibronectin [9, 49]. *PDGFRB* mRNA and PDGFRβ protein expression were observed in angiogenic ECs that formed sprouts [49]. PDGFRβ protein expression was measured at 1900–2900 PDGFRβ/cell on the plasma membrane of HUVECs after 6 hours of coculture with fibroblasts [9]. PDGFRα protein expression was observed in the PECAM^+^ endothelial cells of the outflow tract in mouse embryos [50]. *PDGFRA* deletion in endothelial cells impaired outflow tract development with hypocellular cushions [50]. Minor PDGFRα co-expression was found in CD146^+^ endothelial cells differentiated from PDGFRα^+^ cells isolated from human fetal hearts [51]. PDGFRs also contribute to vasculature development in tumor microenvironments [52]. Approximately 3000 PDGFRα and β per cell were observed on human tumor EC-like cell membranes [14]. PDGFRβ is considered a marker of endothelial-to-mesenchymal transition in the tumor microenvironment and suppresses VEGFR2 expression in the process [53, 54]. ECs secrete PDGF and recruit mural cells to stabilize blood vessels [55, 56]. High expression of PDGFRα and PDGFRβ is associated with high aggressiveness in invasive breast carcinoma and poor survival in prostate cancer stroma, respectively [57, 58]. PDGFRβ protein expression also serves as a therapeutic marker in benign and malignant prostate cancer stroma [57, 58]. Overall, PDGFR expression serves as a pertinent example of the reduced heterogeneity of ECs when monocultured in vitro, reflecting the impact of the isolated environment devoid of organ-specific cues. Our study serves as a robust reference for understanding the abundance of RTKs in monocultured ECs from various tissues.

### ECs in Culture Show Organ-Specific Gene and Protein Expression, Structures, and Functions

Numerous investigations have reported variation in the expression of angiogenesis-related genes in ECs derived from different tissues, in vitro, by measuring the RNA levels, which may be related to these cells’ tissue-specific proliferation, migration, survival, and responses to inflammatory stimuli [59, 60]. Our study reveals, however, that the expression levels of RTK proteins in ECs from dermis, umbilical vein, kidney, liver, and brain are comparable and of consistent magnitudes for each receptor type, both on the plasma membrane and across the whole cell. Previous studies have demonstrated the low correlation between RNA levels, protein expression levels, and functional changes due to translational and post-translational regulation [14, 61–63]. Our findings suggest that the different physiological characteristics of ECs in different organs or tissues may not be significantly attributed to variations in VEGFR, NRP1, or Axl abundance, in vitro.

While cell heterogeneity is a hallmark of site-specific vasculature in vivo, including variation in cell morphology, function, and gene expression [42, 64], our study indicates that monoculture conditions exhibit low heterogeneity in RTK expression, as shown by QE values between 0.2 and 0.7 across all cell types [5, 9, 14]. Skewness corresponds to the degree of symmetry of data distribution and the spread of data around the mean value [65, 66]. Skewness is calculated based on the differences between mode and mean [67]. Positive skewness for each distribution in our study indicated that the majority of the data were concentrated at the lower end of the distribution, with fewer outliners on the higher end [67]. This observation also confirmed the homogeneity of the RTK protein expression pattern of ECs from the same tissue. Kurtosis corresponds to the sharpness of the peak of a distribution [68]. Our analysis showed that kurtosis was larger than 3 for each distribution, indicating a sharper distribution with fewer data points in the tails and suggesting a homogenous expression of each RTK protein in ECs from the same tissue [65]. Notably, variation appeared more prominent on the plasma membrane than across the whole cell, suggesting that protein localization plays a more crucial role than overall expression in defining cellular phenotype and function. This homogeneous RTK density aligns with findings that receptor function remains stable despite changes in concentration by orders of magnitude [69].

Our study reveals some statistically significant differences in RTK abundance between cultured endothelial cells from five organs. However, the observed variability was modest, with the numbers of each receptor being of the same order of magnitude and on the same scale across different cell types (Table 1). Studies have shown that receptor functions are insensitive to volume changes of up to two orders of magnitude [69]. This finding of less complexity in vitro underscores the noteworthy influence of the environment on shaping cellular morphology and functionality. Aird has described endothelial cells as inherently heterogeneous in vivo, even within the same vascular bed, and reported heterogeneity across endothelial cells from various organs [42, 64, 70]. He posited that the environment, including in vitro culture, could induce changes affecting endothelial heterogeneity; however, there may be “site-specific” characteristics innate to ECs that enable retention of those characteristics in vitro [42, 70]. Zheng et al. have also observed that in vitro cell expansion can significantly reduce EC heterogeneity. When they compared freshly isolated heart and kidney ECs, they observed > 5000 differentially expressed genes, whereas after culture, the expression of only 867 genes remained significantly different [59]. It has been suggested that organ-specific EC heterogeneity is regulated by both local availability of growth factors and cell capacities to respond to the signals [42, 71]. Zheng et al. showed differential sprouting capacities in response to VEGF by ECs derived from heart, lung, liver, and kidney, in descending order [59]. Our finding indicates, however, that the differential angiogenic capacities could not be fully captured by RTK quantification in the investigated ECs, in vitro, suggesting a need to check the heterogeneity ex vivo in freshly isolated samples.

ECs originating from various tissues exhibit unique structural and functional characteristics that arise from differences in development and surrounding microenvironments (Table 4) [59, 64]. For instance, dermal ECs show a characteristic cobblestone morphology and play an essential role in tissue repair and wound healing [72]. Due to their robust angiogenic capacities and capacities to produce basement membrane components, dermal ECs are used extensively to create vessel-like structures in vivo and in vitro, often cocultured with fibroblasts in skin tissue engineering [72, 73]. Intriguingly, cultured HDMECs (dermis; *PDGFRA^−/−^* and *PDGFRB^−/−^*) expressed the lowest levels of VEGFR1 and VEGFR2 proteins on the plasma membrane and across the whole cell among the five cell types. This finding underscores the need to examine additional factors that may influence the angiogenic potential of ECs.

ECs from umbilical veins and umbilical arteries display different morphological features, with venous ECs generally exhibiting a thinner profile and shorter and broader shape in comparison with the elongated ellipsoidal appearance of arterial ECs [64, 74, 75]. Other variables regarding cell viability, metabolic activity, membrane integrity, secretion of vasoactive substances, and the production of extracellular matrix proteins are reported to not differ between venous and arterial ECs [74]. In contrast, arteries showed high angiogenic responsiveness and increased sensitivity to angiogenic inhibitors compared with veins [76], suggesting a need to investigate the underlying mechanisms of such differences between veins and arteries.

In the kidney, ECs exhibit relatively small fenestrae and actively synthesize the basement membrane [59, 64]. Kidney-specific inhibition of VEGF has been shown to result in significant phenotypic changes in glomerular ECs and the loss of barrier function [77]. VEGFR3 in glomerular ECs modulates the endothelium’s permeability and regulates the formation of glomerular microvasculature [39, 40]. This aligns with the robust total VEGFR3 protein expression in HGMECs (kidney).

The capillary endothelium of the liver is characterized by its discontinuous nature and possesses large fenestrae, facilitating fluid filtration [78]. We observed relatively high densities of plasma membrane VEGFRs in liver endothelial cells, aligning with the highly proliferative phenotype of liver hepatocytes [79]. Moreover, ECs derived from the liver exhibited higher sprouting capacities than ECs from kidney [59]. Hepatocytes express VEGF, which binds to VEGFR1 and VEGFR2 on endothelial cells, leading to endothelial cell proliferation [79]. As mentioned above, VEGF-C/VEGFR3 signaling regulates sinusoidal vascular growth and is required for normal fetal liver size [41].

Brain endothelium is non-fenestrated and enriched in tight junctions, providing a lower permeability essential for maintaining the integrity of the blood–brain barrier [70, 80]. Elevated glucose levels are essential for nervous tissue function in the brain [81], with brain microvessels facilitating the transport of glucose from the bloodstream into the brain [82, 83]. High glucose concentrations in the endothelium, however, were found to reduce plasma membrane VEGFR2 abundance and impair VEGFR2 activation, while total VEGFR2 protein expression remained unaltered [84]. Without a high glucose microenvironment, the HBMECs (brain) in our study had whole-cell VEGFR2 protein levels comparable to those in ECs from other tissues, in agreement with the existing literature. Notably, the heightened plasma membrane VEGFR2 levels on HBMECs (brain) may be attributed to a phenomenon characterized as “derepression”, the process of removing gene expression inhibition and resulting in increased gene expression.

### Impact of Culture Medium Composition on RTK Protein Expression

Differences in culture media can have significant impacts on RTK expression levels. It is noteworthy that cells from different vendors were cultured with the recommended media containing various components (Table 3). For instance, serum [13] and heparin [92] are two additives that have been found to play a role in regulating receptor levels and cell functions. The presence of high serum in the culture medium has been found to reduce NRP1 protein concentrations on the plasma membranes of human dermal fibroblasts [5]. In our study, HUVECs (umbilical vein) cultivated in low-serum EGM-2 medium with 2% FBS exhibited the highest level of plasma membrane NRP1 protein, corroborating this observation. Moreover, heparin inhibits cell proliferation induced by growth factors [85]. The medium for HDMECs (dermis; *PDGFRA^−/−^* and *PDGFRB^−/−^*) contains a higher concentration of heparin (0.0015 mg/mL) than does the medium for HUVECs (umbilical vein) (0.001 mg/mL). This could potentially contribute to the comparatively lower plasma membrane and total expression levels of VEGFR1 and VEGFR2 proteins in HDMECs (dermis; *PDGFRA^−/−^* and *PDGFRB^−/−^*). However, to conclusively correlate the presence of these additives with the differences that we observed across cells would require further examination of the effects of growth media on these specific cell types.

**Table 3.**
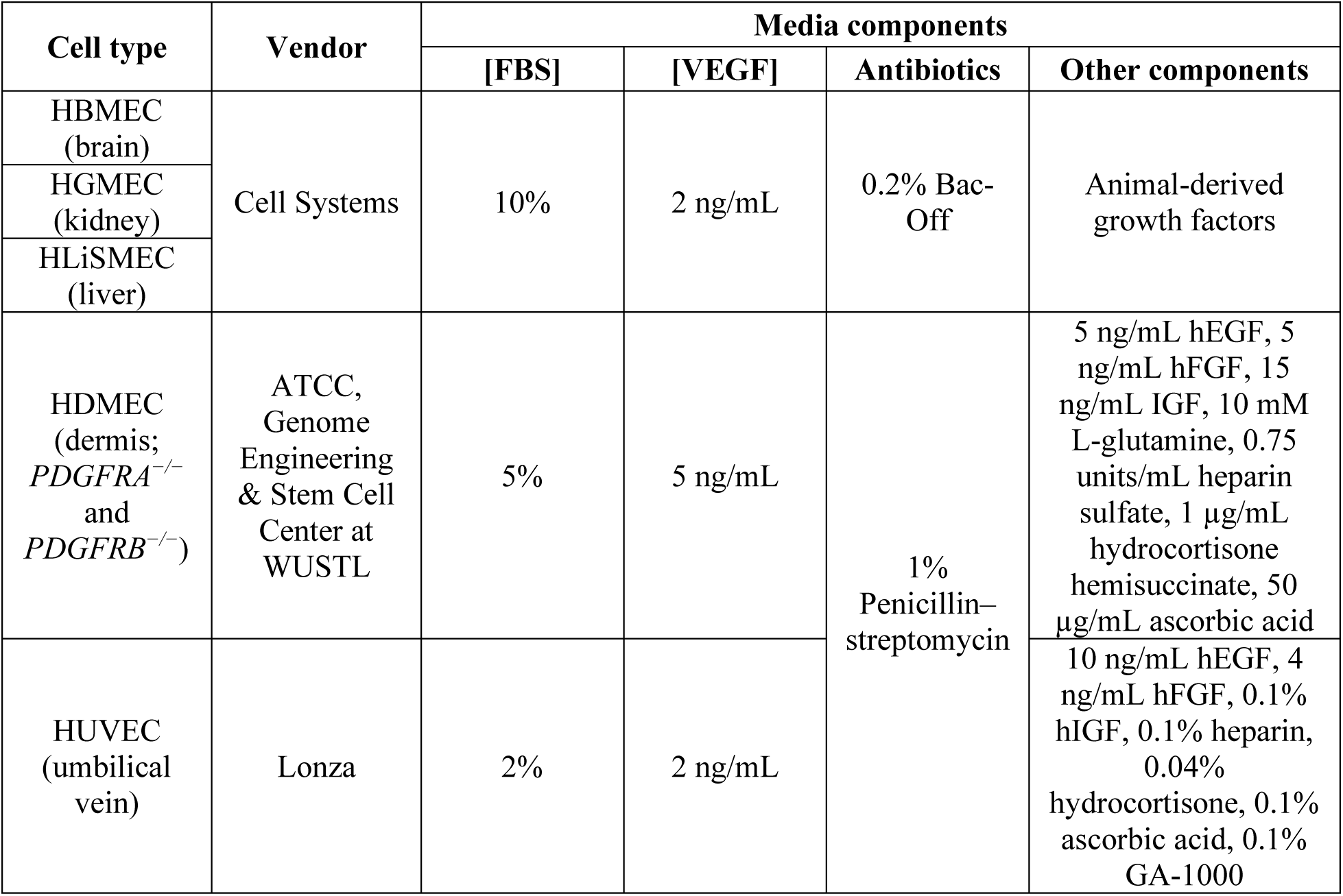
Medium components for each cell type.

**Table 4.** The structural and functional differences among the cell types examined.

| Cell type | Tissue origins | Structural differences | Functional differences |
| --- | --- | --- | --- |
| HDMEC | Dermis in skin | Exhibit cobblestone morphology [72]; high surface area and close to external environment; vasoconstriction and vasodilation control the transfer of body heat [88, 89] | Contribute to tissue repair and wound healing [72]; regulate endothelial permeability, skin homeostasis, and thermoregulation [39, 88, 89] |
| HUVEC | Umbilical vein | Exhibit a thinner profile and shorter and broader shape [74, 75]; the blood vessel wall cannot stretch or retract [90] | Behave as an organ to return the blood flow [90] |
| HGMEC | Kidney glomerulus | Exhibit relatively smaller fenestrae [59, 64] | Actively synthesize the basement membrane [59, 64]; modulate endothelial permeability [40] |
| HLiSMEC | Liver sinusoids | Exhibit gaps and possess large fenestrae [79] | Facilitate fluid filtration, endocytosis [79] |
| HBMEC | Brain | Non-fenestrated and enriched in tight junctions [70, 80] | Lower permeability maintains the integrity of the blood–brain barrier [70, 80] |

### Quantitative receptor study advances computational biology

Quantitative receptor studies have played a pivotal role in the advancement of computational biology, enabling the measurement of cellular heterogeneity and functional capacities [42, 64, 86]. Computational modeling facilitates the simulation of physiological processes by comprehensively integrating all relevant factors, thereby enhancing insight into underlying mechanisms and enabling exploration of potential drugs as well as the determination of optimal drug dosages [42, 64, 86]. Our study makes a notable contribution to this endeavor by offering quantitative measurements of RTK abundance at both the plasma membrane and whole-cell levels. Our findings reveal a remarkable consistency in RTK abundance across cells with five distinct tissue origins, suggesting that comparable RTK levels among different ECs may be assumed, in vitro. Future research should focus on assessing the angiogenic capacities of different EC types through functional assays such as proliferation, migration, and tube formation assays, and elucidating the role of downstream signaling pathways in different cell types.

## Conclusion

Our work enriches the understanding of organ-specificity at both the molecular and cellular levels by detailing RTK distribution patterns across various tissue-derived cells. These insights serve as a foundation for further studies into therapeutic strategies targeting endothelial receptor profiles to modulate angiogenesis effectively.

## Acknowledgments

This material is based on work supported by the National Science Foundation under Grant No. 1923151. Research reported in this publication was also supported by the National Institute of Heart, Lung, and Blood of the National Institutes of Health under Award No. 7R01HL159946-02. Any opinions, findings, conclusions, or recommendations expressed in this material are those of the author(s) and do not necessarily reflect the views of NSF or NIH. Primary human dermal microvascular endothelial cells (HDMECs) (ATCC, CRL-4060) with *PDGFRA* and *PDGFRB* double knock out (DKO) via CRISPR/Cas9 were obtained at the Genome Engineering & Stem Cell Center at Washington University in St. Louis, MO.

## Author Contributions

X.L. and P.I.I. conceived the experiments; X.L. conducted the experiments, analyzed the results, and prepared the figures and tables; X.L., Y.F., and P.I.I. contributed to the writing of the manuscript.

## References

1. Reiterer M, Branco CM (2020) Endothelial cells and organ function: applications and implications of understanding unique and reciprocal remodelling. FEBS J 287:1088–1100

2. Urbanczyk M, Zbinden A, Schenke-Layland K (2022) Organ-specific endothelial cell heterogenicity and its impact on regenerative medicine and biomedical engineering applications. Adv Drug Deliv Rev 186:114323

3. Jeltsch M, Leppanen V-M, Saharinen P, Alitalo K (2013) Receptor tyrosine kinase-mediated angiogenesis. Cold Spring Harb Perspect Biol 5:a009183–a009183

4. Virág J, Kenessey I, Haberler C, Piurkó V, Bálint K, Döme B, T\’\imár J, Garami M, Hegedűs B (2014) Angiogenesis and angiogenic tyrosine kinase receptor expression in pediatric brain tumors. Pathol Oncol Res 20:417–426

5. Chen S, Guo X, Imarenezor O, Imoukhuede PI (2015) Quantification of VEGFRs, NRP1, and PDGFRs on Endothelial Cells and Fibroblasts Reveals Serum, Intra-Family Ligand, and Cross-Family Ligand Regulation. Cell Mol Bioeng 8:. 10.1007/s12195-015-0411-x

6. Imoukhuede PI, Popel AS (2012) Expression of VEGF receptors on endothelial cells in mouse skeletal muscle. PLoS One 7:e44791

7. Chen S, Weddell J, Gupta P, Conard G, Parkin J, Imoukhuede PI (2017) qFlow Cytometry-Based Receptoromic Screening: A High-Throughput Quantification Approach Informing Biomarker Selection and Nanosensor Development

8. Imoukhuede PI, Popel AS (2011) Quantification and cell-to-cell variation of vascular endothelial growth factor receptors. Exp Cell Res 317:955–965

9. Chen S, Imoukhuede PI (2019) Single-cell receptor quantification of an in vitro coculture angiogenesis model reveals VEGFR, NRP1, Tie2, and PDGFR regulation and endothelial heterogeneity. Processes (Basel) 7:356

10. Fang Y, Malik M, England SK, Imoukhuede PI (2022) Absolute quantification of plasma membrane receptors via quantitative flow cytometry. In: Methods in Molecular Biology. Springer US, New York, NY, pp 61–77

11. Imoukhuede PI, Popel AS (2014) Quantitative fluorescent profiling ofVEGFRs reveals tumor cell and endothelial cell heterogeneity in breast cancer xenografts. Cancer Med 3:225–244

12. Fang Y, Chen L, Imoukhuede PI (2023) Toward blood-based precision medicine: Identifying age-sex-specific vascular biomarker quantities on circulating vascular cells. Cell Mol Bioeng 16:189–204

13. Fang Y, Imoukhuede PI (2023) Axl and vascular endothelial growth factor receptors exhibit variations in membrane localization and heterogeneity across monolayer and spheroid high-grade serous ovarian cancer models. GEN Biotechnol 2:43–56

14. Chen S, Le T, Harley BAC, Imoukhuede PI (2018) Characterizing glioblastoma heterogeneity via single-cell receptor quantification. Front Bioeng Biotechnol 6:

15. Weddell JC, Imoukhuede PI (2014) Quantitative characterization of cellular membrane-receptor heterogeneity through statistical and computational modeling. PLoS One 9:e97271

16. Imoukhuede PI, Dokun AO, Annex BH, Popel AS (2013) Endothelial cell-by-cell profiling reveals the temporal dynamics of VEGFR1 and VEGFR2 membrane localization after murine hindlimb ischemia. Am J Physiol Heart Circ Physiol 304:H1085–H1093

17. Weddell JC, Imoukhuede PI (2018) Computational systems biology for the VEGF family in angiogenesis. In: Encyclopedia of Cardiovascular Research and Medicine. Elsevier, pp 659–676

18. Nakayama M, Berger P (2013) Coordination of VEGF receptor trafficking and signaling by coreceptors. Exp Cell Res 319:1340–1347

19. Wiszniak S, Schwarz Q (2021) Exploring the intracrine functions of VEGF-A. Biomolecules 11:128

20. Weddell JC, Imoukhuede PI (2017) Integrative meta-modeling identifies endocytic vesicles, late endosome and the nucleus as the cellular compartments primarily directing RTK signaling. Integr Biol (Camb) 9:464–484

21. Fang Y, Reinl EL, Liu A, Prochaska TD, Malik M, Frolova AI, England SK, Imoukhuede PI (2024) Quantification of surface-localized and total oxytocin receptor in myometrial smooth muscle cells. Heliyon 10:e25761

22. Malik M, Ward MD, Fang Y, Porter JR, Zimmerman MI, Koelblen T, Roh M, Frolova AI, Burris TP, Bowman GR, Imoukhuede PI, England SK (2021) Naturally occurring genetic variants in the oxytocin receptor alter receptor signaling profiles. ACS Pharmacol Transl Sci 4:1543–1555

23. Rao CR (1982) Diversity and dissimilarity coefficients: A unified approach. Theor Popul Biol 21:24–43

24. Pavoine S, Dolédec S (2005) The apportionment of quadratic entropy: a useful alternative for partitioning diversity in ecological data. Environ Ecol Stat 12:125–138

25. Botta-Dukát Z (2005) Rao’s quadratic entropy as a measure of functional diversity based on multiple traits. J Veg Sci 16:533–540

26. Roy S, Bag AK, Singh RK, Talmadge JE, Batra SK, Datta K (2017) Multifaceted role of neuropilins in the immune system: Potential targets for immunotherapy. Front Immunol 8:

27. Wittko-Schneider IM, Schneider FT, Plate KH (2013) Brain homeostasis: VEGF receptor 1 and 2— two unequal brothers in mind. Cell Mol Life Sci 70:1705–1725

28. Carmeliet P, Moons L, Luttun A, Vincenti V, Compernolle V, De Mol M, Wu Y, Bono F, Devy L, Beck H, Scholz D, Acker T, DiPalma T, Dewerchin M, Noel A, Stalmans I, Barra A, Blacher S, Vandendriessche Thierry and Ponten A, Eriksson U, Plate KH, Foidart J-M, Schaper W, Charnock-Jones DS, Hicklin DJ, Herbert J-M, Collen D, Persico MG (2001) Synergism between vascular endothelial growth factor and placental growth factor contributes to angiogenesis and plasma extravasation in pathological conditions. Nat Med 7:575–583

29. Waltenberger J, Claesson-Welsh L, Siegbahn A, Shibuya M, Heldin CH (1994) Different signal transduction properties of KDR and Flt1, two receptors for vascular endothelial growth factor. J Biol Chem 269:26988–26995

30. Sawano A, Iwai S, Sakurai Y, Ito M, Shitara K, Nakahata T, Shibuya M (2001) Flt-1, vascular endothelial growth factor receptor 1, is a novel cell surface marker for the lineage of monocyte-macrophages in humans. Blood 97:785–791

31. Ogawa S, Oku A, Sawano A, Yamaguchi S, Yazaki Y, Shibuya M (1998) A novel type of vascular endothelial growth factor, VEGF-E (NZ-7 VEGF), preferentially utilizes KDR/flk-1 receptor and carries a potent mitotic activity without heparin-binding domain. J Biol Chem 273:31273–31282

32. Shalaby F, Rossant J, Yamaguchi TP, Gertsenstein M, Wu X-F, Breitman ML, Schuh AC (1995) Failure of blood-island formation and vasculogenesis in Flk-1-deficient mice. Nature 376:62–66

33. Roberts DM, Kearney JB, Johnson JH, Rosenberg MP, Kumar R, Bautch VL The Vascular Endothelial Growth Factor (VEGF) Receptor Flt-1 (VEGFR-1) Modulates Flk-1 (VEGFR-2) Signaling During Blood Vessel Formation

34. Sarabipour S, Kinghorn K, Quigley Kaitlyn M and Kovacs-Kasa A, Annex BH, Bautch Victoria L and Mac Gabhann F (2022) Trafficking dynamics of VEGFR1, VEGFR2, and NRP1 in human endothelial cells

35. Sarabipour S, Kinghorn K, Quigley Kaitlyn M and Kovacs-Kasa A, Annex BH, Bautch Victoria L and Mac Gabhann F (2024) Impact of ligand binding on VEGFR1, VEGFR2, and NRP1 localization in human endothelial cells

36. Napione L, Pavan S, Veglio A, Picco A, Boffetta G, Celani A, Seano Giorgio and Primo L, Gamba A, Bussolino F (2012) Unraveling the influence of endothelial cell density on VEGF-A signaling. Blood 119:5599–5607

37. Hagura A, Asai J, Maruyama K, Takenaka H, Kinoshita S, Katoh N (2014) The VEGF-C/VEGFR3 signaling pathway contributes to resolving chronic skin inflammation by activating lymphatic vessel function. J Dermatol Sci 73:135–141

38. Deng Y, Zhang X, Simons M (2015) Molecular controls of lymphatic VEGFR3 signaling. Arterioscler Thromb Vasc Biol 35:421–429

39. Heinolainen K, Karaman S, D’Amico G, Tammela T, Sormunen R, Eklund L, Alitalo K, Zarkada G (2017) VEGFR3 modulates vascular permeability by controlling VEGF/VEGFR2 signaling. Circ Res 120:1414–1425

40. Donnan MD, Deb DK, Onay T, Scott RP, Ni E, Zhou Y, Quaggin SE (2023) Formation of the glomerular microvasculature is regulated by VEGFR-3. Am J Physiol Renal Physiol 324:F91–F105

41. Sung DC, Chen M, Dominguez MH, Mahadevan A, Chen X, Yang J, Gao S, Ren AA, Tang AT, Mericko P, Patton R, Lee M, Jannaway M, Nottebaum AF, Vestweber D, Scallan Joshua P and Kahn ML (2022) Sinusoidal and lymphatic vessel growth is controlled by reciprocal VEGF-C–CDH5 inhibition. Nat Cardiovasc Res 1:1006–1021

42. Aird WC (2012) Endothelial cell heterogeneity. Cold Spring Harb Perspect Med 2:a006429–a006429

43. Shintani Y, Takashima S, Asano Y, Kato H, Liao Y, Yamazaki S, Tsukamoto O, Seguchi O, Yamamoto H, Fukushima T, Sugahara K, Kitakaze M, Hori M (2006) Glycosaminoglycan modification of neuropilin-1 modulates VEGFR2 signaling. EMBO J 25:3045–3055

44. Gabhann F Mac, Popel AS (2006) Targeting neuropilin-1 to inhibit VEGF signaling in cancer: Comparison of therapeutic approaches. PLoS Comput Biol 2:e180

45. Bosseboeuf E, Chikh A, Chaker AB, Mitchell TP, Vignaraja D, Rajendrakumar R, Khambata RS, Nightingale TD, Mason JC, Randi AM, Ahluwalia A, Raimondi C (2023) Neuropilin-1 interacts with VE-cadherin and TGFBR2 to stabilize adherens junctions and prevent activation of endothelium under flow. Sci Signal 16:

46. Gioelli N, Neilson LJ, Wei N, Villari G, Chen W, Kuhle B, Ehling Manuel and Maione F, Willox S, Brundu S, Avanzato D, Koulouras G, Mazzone M, Giraudo E, Yang X-L, Valdembri D, Zanivan S, Serini G (2022) Neuropilin 1 and its inhibitory ligand mini-tryptophanyl-tRNA synthetase inversely regulate VE-cadherin turnover and vascular permeability. Nat Commun 13:

47. Shao H, Teramae D, Wells A (2023) Axl contributes to efficient migration and invasion of melanoma cells. PLoS One 18:e0283749

48. Konno K, Yamasaki M, Miyazaki T, Watanabe M (2023) Glyoxal fixation: An approach to solve immunohistochemical problem in neuroscience research. Sci Adv 9:

49. Battegay EJ, Rupp J, Iruela-Arispe L, Sage EH, Pech M (1994) PDGF-BB modulates endothelial proliferation and angiogenesis in vitro via PDGF beta-receptors. J Cell Biol 125:917–928

50. Aghajanian H, Cho YK, Rizer NW, Wang Q, Li L, Degenhardt K, Jain R (2017) Pdgfrα functions in endothelial-derived cells to regulate neural crest cells and development of the great arteries. Dis Model Mech

51. Chong JJH, Reinecke H, Iwata M, Torok-Storb B, Stempien-Otero A, Murry CE (2013) Progenitor cells identified by PDGFR-alpha expression in the developing and diseased human heart. Stem Cells Dev 22:1932–1943

52. Andrae J, Gallini R, Betsholtz C (2008) Role of platelet-derived growth factors in physiology and medicine. Genes Dev 22:1276–1312

53. Valero-Muñoz M, Oh A, Faudoa Elizabeth and Bretón-Romero R, El Adili F, Bujor A, Sam F (2021) Endothelial-mesenchymal transition in heart failure with a preserved ejection fraction: Insights into the cardiorenal syndrome. Circ Heart Fail 14:

54. Yin Z, Wang L (2023) Endothelial-to-mesenchymal transition in tumour progression and its potential roles in tumour therapy. Ann Med 55:1058–1069

55. Hellström M, Kalén M, Lindahl P, Abramsson A, Betsholtz C (1999) Role of PDGF-B and PDGFR-β in recruitment of vascular smooth muscle cells and pericytes during embryonic blood vessel formation in the mouse. Development 126:3047–3055

56. Lindahl P (1997) Pericyte Loss and Microaneurysm Formation in PDGF-B-Deficient Mice. Science (1979) 277:. 10.1126/science.277.5323.242

57. Carvalho I, Milanezi F, Martins A, Reis RM, Schmitt F (2005) Overexpression of platelet-derived growth factor receptor α in breast cancer is associated with tumour progression. Breast Cancer Res 7:

58. Nordby Y, Richardsen E, Rakaee M, Ness N, Donnem T, Patel HRH, Busund Lill-Tove and Bremnes RM, Andersen S (2017) High expression of PDGFR-β in prostate cancer stroma is independently associated with clinical and biochemical prostate cancer recurrence. Sci Rep 7:

59. Marcu R, Choi YJ, Xue J, Fortin CL, Wang Y, Nagao RJ, Xu J, MacDonald JW, Bammler TK, Murry CE, Muczynski K, Stevens KR, Himmelfarb Jonathan and Schwartz SM, Zheng Y (2018) Human organ-specific endothelial cell heterogeneity. iScience 4:20–35

60. Chi J-T, Chang HY, Haraldsen G, Jahnsen FL, Troyanskaya OG, Chang DS, Wang Z, Rockson SG, van de Rijn M, Botstein D, Brown PO (2003) Endothelial cell diversity revealed by global expression profiling. Proc Natl Acad Sci U S A 100:10623–10628

61. Nie L, Wu G, Zhang W (2006) Correlation of mRNA expression and protein abundance affected by multiple sequence features related to translational efficiency in Desulfovibrio vulgaris: A quantitative analysis. Genetics 174:2229–2243

62. Greenbaum D, Colangelo C, Williams Kenneth and Gerstein M (2003) Comparing protein abundance and mRNA expression levels on a genomic scale. Genome Biol 4:117

63. Beyer A, Hollunder J, Nasheuer H-P, Wilhelm T (2004) Post-transcriptional expression regulation in the yeast Saccharomyces cerevisiae on a genomic scale. Mol Cell Proteomics 3:1083–1092

64. Aird WC (2007) Phenotypic heterogeneity of the endothelium: II. Representative vascular beds. Circ Res 100:174–190

65. Folcarelli R, van Staveren S, Tinnevelt Gerjen and Cadot E, Vrisekoop N, Buydens L, Koenderman L, Jansen J, van den Brink OF (2022) Transformation of multicolour flow cytometry data with OTflow prevents misleading multivariate analysis results and incorrect immunological conclusions. Cytometry A 101:72–85

66. D’agostino RB, Belanger A, D’agostino Jr RB (1990) A suggestion for using powerful and informative tests of normality. Am Stat 44:316–321

67. Heins A-L, Johanson T, Han S, Lundin L, Carlquist M, Gernaey K V, Sørensen SJ, Eliasson Lantz A (2019) Quantitative flow cytometry to understand population heterogeneity in response to changes in substrate availability in Escherichia coli and Saccharomyces cerevisiae chemostats. Front Bioeng Biotechnol 7:

68. Balanda KP, Macgillivray HL (1988) Kurtosis: A critical review. Am Stat 42:111–119

69. Shankaran H, Resat H, Wiley HS (2007) Cell surface receptors for signal transduction and ligand transport: A design principles study. PLoS Comput Biol 3:e101

70. Aird WC (2005) Spatial and temporal dynamics of the endothelium. J Thromb Haemost 3:1392–1406

71. Préau L, Lischke A, Merkel M, Oegel N, Weissenbruch M, Michael A, Park H, Gradl D, Kupatt C, le Noble F (2024) Parenchymal cues define Vegfa-driven venous angiogenesis by activating a sprouting competent venous endothelial subtype. Nat Commun 15:

72. Zeng Q, Macri LK, Prasad A, Clark RAF, Zeugolis DI, Hanley C, Garcia Y, Pandit A, Leavesley DI, Stupar D, Fernandez ML, Fan C, Upton Z (2017) 6.20 skin tissue engineering ☆. In: Comprehensive Biomaterials II. Elsevier, pp 334–382

73. Bourland J, Fradette J (2018) Strategies to promote the vascularization of skin substitutes after transplantation. In: Skin Tissue Models for Regenerative Medicine. Elsevier, pp 177–200

74. Lau S, Gossen M, Lendlein A, Jung F (2021) Venous and arterial endothelial cells from human umbilical cords: Potential cell sources for cardiovascular research. Int J Mol Sci 22:978

75. dela Paz NG, D’Amore PA (2009) Arterial versus venous endothelial cells. Cell Tissue Res 335:5–16

76. Blebea J, Vu J-H, Assadnia S, McLaughlin PJ, Atnip RG, Zagon IS (2002) Differential effects of vascular growth factors on arterial and venous angiogenesis. J Vasc Surg 35:532–538

77. Eremina V, Sood M, Haigh J, Nagy A, Lajoie G, Ferrara N, Gerber H-P, Kikkawa Y, Miner JH, Quaggin SE (2003) Glomerular-specific alterations of VEGF-A expression lead to distinct congenital and acquired renal diseases. J Clin Invest 111:707–716

78. Braet F, Wisse E (2002) Structural and functional aspects of liver sinusoidal endothelial cell fenestrae: a review. Comp Hepatol 1:

79. LeCouter J, Moritz DR, Li B, Phillips GL, Liang XH, Gerber H-P, Hillan KJ, Ferrara N (2003) Angiogenesis-independent endothelial protection of liver: Role of VEGFR-1. Science (1979) 299:890–893

80. Nitta T, Hata M, Gotoh S, Seo Y, Sasaki H, Hashimoto N, Furuse M, Tsukita S (2003) Size-selective loosening of the blood-brain barrier in claudin-5–deficient mice. J Cell Biol 161:653–660

81. Joo F (1996) Endothelial cells of the brain and other organ systems: Some similarities and differences. Prog Neurobiol 48:255–273

82. Goldstein GW (1979) Relation of potassium transport to oxidative metabolism in isolated brain capillaries. J Physiol 286:185–195

83. Mrsulja BB, Djuricic BM, Micic DF (1980) Circulatory and Developmental Aspects of Brain Metabolism. Plenum Press, New York

84. Warren CM, Ziyad S, Briot A, Der A, Iruela-Arispe ML (2014) A ligand-independent VEGFR2 signaling pathway limits angiogenic responses in diabetes. Sci Signal 7:

85. Herrick WG, Rattan S, Nguyen T V, Grunwald MS, Barney CW, Crosby AJ, Peyton SR (2015) Smooth muscle stiffness sensitivity is driven by soluble and insoluble ECM chemistry. Cell Mol Bioeng 8:333–348

86. Mojiri A, Alavi P, Lorenzana Carrillo MA, Nakhaei-Nejad M, Sergi CM, Thebaud B, Aird WC, Jahroudi N (2019) Endothelial cells of different organs exhibit heterogeneity in von Willebrand factor expression in response to hypoxia. Atherosclerosis 282:1–10

87. Lee S, Mandic J, Van Vliet KJ (2007) Chemomechanical mapping of ligand–receptor binding kinetics on cells. Proc Natl Acad Sci U S A 104:9609–9614

88. Gutterman DD, Chabowski DS, Kadlec Andrew O and Durand MJ, Freed JK, Ait-Aissa Karima and Beyer AM (2016) The human microcirculation: Regulation of flow and beyond: Regulation of flow and beyond. Circ Res 118:157–172

89. Charkoudian N (2003) Skin blood flow in adult human thermoregulation: how it works, when it does not, and why. Mayo Clin Proc 78:603–612

90. Hoang-Ngoc Minh, Gebrane-Younes J, Smadja A, Orcel L (1985) Structure and function of the fetal umbilical vessels. J Gynecol Obstet Biol Reprod (Paris) 14:973–979

