## Supplementary material for "Cultured endothelial cells present organ-specific RTK distributions: advancing receptor measurement and data standardization via quantitative flow cytometry": Suppl. Data

### Supplementary Data

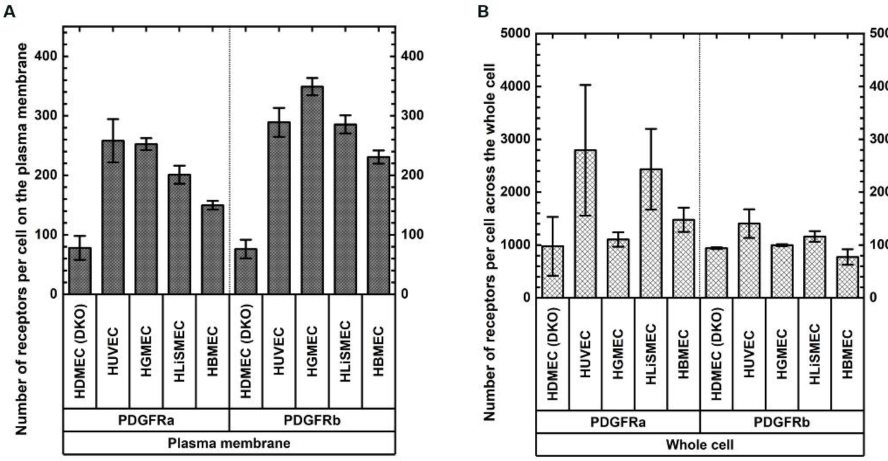


**Fig. S1 Monocultured ECs expressed few to no PDGFRα or β on the plasma membrane (A) and low levels across the whole cell (B).** The efficiency of DKO of HDMECs (dermis; *PDGFRA*$^{-/-}$ and *PDGFRB*$^{-/-}$) was validated by next-generation sequencing (NGS) analysis (data not shown). The numbers of PDGFRs resulted from nonspecific binding between PDGFR antibodies and PDGFRs.

**Table S1 NGS validation was employed to assess the efficiency of PDGFRα and β double knockout.** The gRNAs, YM200.sp17 and YM199.sp7, were utilized to target PDGFRα and PDGFRβ, respectively. Subsequent NGS analysis was conducted to confirm the knockout efficiency in both single KO and double KO clones. Specifically, the D11.DKO and E4.DKO cell lines were assessed for both target sites. The E4.DKO cell line was selected for further study because of its lower frequency of insertion–deletion mutations (indels) beyond the exon region.

| YM200-hPDGFRA | HDMECs | Total | sp17 | #1-Indel | #1-Reads(%) | #2-Indel | #2-Reads(%) | #3-Indel | #3-Reads(%) | #4-Indel | #4-Reads(%) |
|---|---|---|---|---|---|---|---|---|---|---|---|
| GEIC-Plate28-B01 | WT | 2486 | 2451 (98.6%) | 0 | 2468 (99.3%) | -1 | 18 (0.7%) | NA | | NA | |
| GEIC-Plate28-B02 | D5- YM199 | no reads | | | | | | | | | |
| GEIC-Plate28-B03 | H9- YM199 | no reads | | | | | | | | | |
| GEIC-Plate28-B04 | B4- YM200 | 3859 | 0 (0.0%) | -10 | 3847 (99.7%) | -11 | 11 (0.3%) | -73 | 1 (0.0%) | NA | |
| GEIC-Plate28-B05 | E4- YM200 | 3727 | 0 (0.0%) | 2 | 1922 (51.6%) | -8 | 1787 (47.9%) | -9 | 10 (0.3%) | 1 | 8 (0.2%) |
| GEIC-Plate28-B06 | D11.DKO | 3652 | 0 (0.0%) | -32 | 1999 (54.7%) | -8 | 1638 (44.9%) | -9 | 7 (0.2%) | -33 | 4 (0.1%) |
| GEIC-Plate28-B07 | E4.DKO | 3941 | 0 (0.0%) | -8 | 1996 (50.6%) | -5 | 1925 (48.8%) | -9 | 9 (0.2%) | -6 | 9 (0.2%) |

| YM199-hPDGFRB | HDMECs | Total | sp7 | #1-Indel | #1-Reads(%) | #2-Indel | #2-Reads(%) | #3-Indel | #3-Reads(%) | #4-Indel | #4-Reads(%) |
|---|---|---|---|---|---|---|---|---|---|---|---|
| GEIC-Plate28-A01 | WT | 2609 | 2569 (98.5%) | 0 | 2595 (99.5%) | -1 | 13 (0.5%) | 1 | 1 (0.0%) | NA | |
| GEIC-Plate28-A02 | D5- YM199 | 2864 | 0 (0.0%) | -7 | 1850 (64.6%) | -23 | 1006 (35.1%) | -8 | 4 (0.1%) | -24 | 2 (0.1%) |
| GEIC-Plate28-A03 | H9- YM199 | 2622 | 0 (0.0%) | 1 | 2615 (99.7%) | 0 | 7 (0.3%) | NA | | NA | |
| GEIC-Plate28-A04 | B4- YM200 | 3033 | 2987 (98.5%) | 0 | 3025 (99.7%) | -1 | 6 (0.2%) | 1 | 1 (0.0%) | -2 | 1 (0.0%) |
| GEIC-Plate28-A05 | E4- YM200 | 2788 | 0 (0.0%) | -5 | 1160 (41.6%) | -6 | 1074 (38.5%) | 2 | 278 (10.0%) | -7 | 274 (9.8%) |
| GEIC-Plate28-A06 | D11.DKO | 3315 | 0 (0.0%) | -11 | 1699 (51.3%) | -1 | 1599 (48.2%) | -12 | 7 (0.2%) | -2 | 6 (0.2%) |
| GEIC-Plate28-A07 | E4.DKO | 3394 | 0 (0.0%) | -1 | 1667 (49.1%) | -25 | 1244 (36.7%) | -10 | 469 (13.8%) | -2 | 8 (0.2%) |

**Table S2 Summary of RTK quantification (molecules/cell) through qFlow, radio-, and biotin labeling in vitro**

| | **Plasma membrane** | | | **Whole cell** | |
|---|---|---|---|---|---|
| | **HDMEC (qFlow)** | **HUVEC** | | | |
| | | **qFlow** | **Radiolabeling** | **Radiolabeling** | **Biotin labeling** |

| | | | | | | | | | |
|---|---|---|---|---|---|---|---|---|---|
| **VEGFR1** | ~1900 [8] | ~1800 [8] | 990 ± 50 [5] | ~1400 [7] | 1800 ± 100 [6] | | | | 19,813 [34] |
| **VEGFR2** | ~7400 [8] | ~4900 [8] | 1890 ± 110 [5] | ~1850 [7] | 5800 ± 300 [6] | 390,000 [36] | 147,000 ± 38,000 [87] | 640,000 [36] | 9661 [34] |
| **VEGFR3** | ~5200 [8] | ~2800 [8] | 1840 ± 200 [5] | ~1700 [7] | | | | | |
| **NRP1** | ~66,000 [8] | ~68,000 [8] | 44,090 ± 330 [5] | ~42,000 [7] | | | | | 98,167 [34] |

17
